# Computational Neuroimaging of Cognition-Emotion Interactions: Affective and Task-similar Interference Differentially Impact Working Memory

**DOI:** 10.1101/2021.06.03.446811

**Authors:** Jie Lisa Ji, Grega Repovs, Genevieve J. Yang, Aleksandar Savic, John D. Murray, Alan Anticevic

**Affiliations:** Department of Psychiatry, Yale University School of Medicine, 300 George Street, New Haven, CT 06511, USA; Department of Psychology, University of Ljubljana, 1000 Ljubljana, Slovenia; Department of Psychiatry, Mount Sinai School of Medicine, 1 Gustave L. Levy Pl, New York, NY 10029; University Psychiatric Hospital Vrapce, University of Zagreb, Zagreb 10000, Croatia; Department of Physics, Yale University, 217 Prospect St, New Haven, CT 06520, USA; Abraham Ribicoff Research Facilities, Connecticut Mental Health Center, New Haven, CT 06519, USA; Department of Psychology, Yale University, 2 Hillhouse Avenue, CT 06520, USA

**Keywords:** working memory, distraction, cognition, emotion, computational modeling, amygdala, functional connectivity

## Abstract

Cognition depends on resisting interference and responding to relevant stimuli. Distracting information, however, varies based on content, requiring distinct filtering mechanisms. For instance, affective information captures attention, disrupts performance and attenuates activation along frontal-parietal regions during cognitive engagement, while recruiting bottom-up regions. Conversely, distraction matching task features (i.e. task-similar) increases fronto-parietal activity. Neural mechanisms behind unique effects of different distraction on cognition remain unknown. Using fMRI in 45 adults, we tested whether affective versus task-similar interference show distinct signals during delayed working memory (WM). We found robust differences between distractor types along fronto-parietal versus affective-ventral neural systems. We studied a hypothesized mechanism of this effect via a biophysically-based computational WM model that implements a functional antagonism between affective/cognitive neural ‘modules’. This architecture reproduced experimental effects: task-similar distractors increased, whereas affective distractors attenuated cognitive module activity while increasing affective module signals. The model architecture suggested that task-based connectivity may be altered in affective-ventral vs. fronto-parietal networks depending on distractor type. Empirically, affective interference significantly increased connectivity within the affective-ventral network, but reduced connectivity between affective-ventral and fronto-parietal networks, which predicted WM performance. These findings detail an antagonistic architecture between cognitive and affective systems, capable of flexibly engaging distinct distractions during cognition.

## INTRODUCTION

The brain encompasses systems that evolved to perform distinct computations, which at times compete in the service of adaptive behavior. Two computations that, at the behavioral and neural level, represent ‘antagonistic’ interactions are affect and cognition, capturing the intuition that logic and reason are at odds with heightened affect (Pessoa, 2008). Affective computations (Lang and Davis, 2006; LeDoux, 2000; Ohman, 2005; Vuilleumier, 2005) allow rapid action deployment (Lang and Davis, 2006; LeDoux, 2000; Ohman, 2005; Vuilleumier, 2005) with privileged access to neural resources (Morris et al., 1998; Ohman et al., 2001; Pessoa, 2005; Vuilleumier and Driver, 2007), at times disrupting cognitive operations (Dolcos and McCarthy, 2006). However, some incoming information may share features with ongoing cognitive processes, and should be integrated, or filtered, using distinct neural mechanisms (Dolcos et al., 2008) – a function hypothesized to rely on ‘top-down’ cognitive control (Cole et al., 2013; Miller and Cohen, 2001). The neural mechanisms behind dissociable effects of distinct distraction on cognition remain uncharacterized.

One canonical cognitive process is working memory (WM) (Baddeley and Hitch, 1974), supported by a network of regions, including fronto-parietal areas (Curtis et al., 2004). WM provides a strong test-bed for understanding effects of distraction on cognition for two reasons: first, its neural architecture is well understood through primate neurophysiology (Funahashi et al., 1989), computational modeling (Compte et al., 2000; Wang, 2010) and human neuroimaging (Curtis et al., 2004; Wager and Smith, 2003; Wager et al., 2014); second, computational models that detail its cellular-level functional architecture can be expanded to generate testable system-level neuroimaging predictions (Anticevic et al., 2012b).

Prior work demonstrated a dichotomy between fronto-parietal and ventral-affective areas when affective distraction appeared during the delay period of WM (Dolcos et al., 2008): Fronto-parietal regions’ activity attenuated, whereas ventral regions exhibited increased activity in response to affective distractors. This may reflect fronto-parietal regions being driven ‘off-line’ by regions processing affective interference, such as the amygdala. Conversely, distractors that matched task features (i.e. task-similar) exerted the opposite effect, increasing fronto-parietal signals (Anticevic et al., 2010a; Anticevic et al., 2011), perhaps requiring distinct ‘filtering’ mechanisms.

Using functional neuroimaging, we first tested for a whole-brain functional segregation between affective and task-similar interference during WM. Building on prior work, we hypothesized that, during delayed WM, ventral systems would exhibit higher signal following affective than task-similar interference. Conversely, we hypothesized that task-similar distraction would be associated with elevated fronto-parietal signal, whereas affective distraction would attenuate these signals. Next, to study possible circuit-level mechanisms behind these phenomena, we used a biophysically-based computational model of WM (Murray et al., 2014) extended to neural systems (Anticevic et al., 2012b), implementing a functional antagonism between affective and cognitive modules (Drevets and Raichle, 1998; Mayberg et al., 1999). The model provided a parsimonious mechanism for observed responses, and made dissociable predictions for the neural and behavioral impact of bi-directional net-inhibitory projections between affective and cognitive systems. Furthermore, the model architecture suggested that task-based connectivity is altered in ventral-affective vs. fronto-parietal areas depending on distractor type, which we tested and related to behavioral performance. These neuroimaging and computational effects detail a proposed antagonistic architecture between cognitive and affective systems, capable of flexibly engaging distinct distractors during cognition with implication for neuropsychiatric conditions where such computations may be disrupted.

## MATERIALS AND METHODS

### Subjects

45 neurologically and psychiatrically healthy right-handed, healthy adults (25 male, mean age=31.2) were recruited from Washington University in Saint Louis and the general community via advertisement by the Psychology Department subject coordinator and underwent neuroimaging data collection.

### Ethics Statement

The study was conducted in accordance with the Declaration of Helsinki. All participants completed and signed an informed consent approved by the Washington University in Saint Louis institutional review broad (IRB) and were paid $25 an hour for their participation. All secondary analyses reported here were approved by the Yale IRB.

### Materials

All participants performed a modified WM delayed response task (Sternberg, 1969) with three distractor types presented during the delay period: i) affectively negative image, ii) visually complex neutral image; and iii) task-related geometric shape. Some aspects of our experiment were reported previously (Anticevic et al., 2011; Anticevic et al., 2010b). Here we studied a key novel and independent question: we specifically characterized the opposing effects of task-similar and affective interference during WM delay at the whole-brain level to better understand the large-scale functional architecture behind opposing distractor effects, previously reported in specific prefrontal/parietal areas (Dolcos et al., 2008). We did so because prior work showed that task-similar distraction (i.e. distractors sharing task properties) was associated with *increased* signals in some dorsal cortical regions rather than the *decreased* signals found for affectively negative distraction (Dolcos et al., 2008). We also examined trials without distraction, used here to estimate distractor-free delay activity as well as trials involving neutral distractors, to establish whole-brain specificity of task-similar versus affectively negative interference. As in our prior studies, the WM sets and task-related distractors were constructed from complex geometric shapes (Attneave and Arnoult, 1956) that were difficult to verbally encode and were generated using an automated Matlab algorithm (Collin and McMullen, 2002) (see (Anticevic et al., 2010b) for more detail on stimulus generation). Negative and neutral visual distractors were selected from the extensively validated International Affective Picture System (IAPS) stimulus set (Lang et al., 1999) and were equated on luminance, contrast, figure-ground relationships, spatial frequency and color (Bradley et al., 2007; Delplanque et al., 2007; Sabatinelli et al., 2005). All distractors were presented centrally, subtending a visual angle of 8.5 degrees. In this study we focused explicitly on the brain-wide signals modulated differentially by affectively negative vs. task-similar distractors (see **Figure 1**). As described below, neutral images were used only to verify specificity of distractor effects (see **Figure 3**).

**Figure 1.**
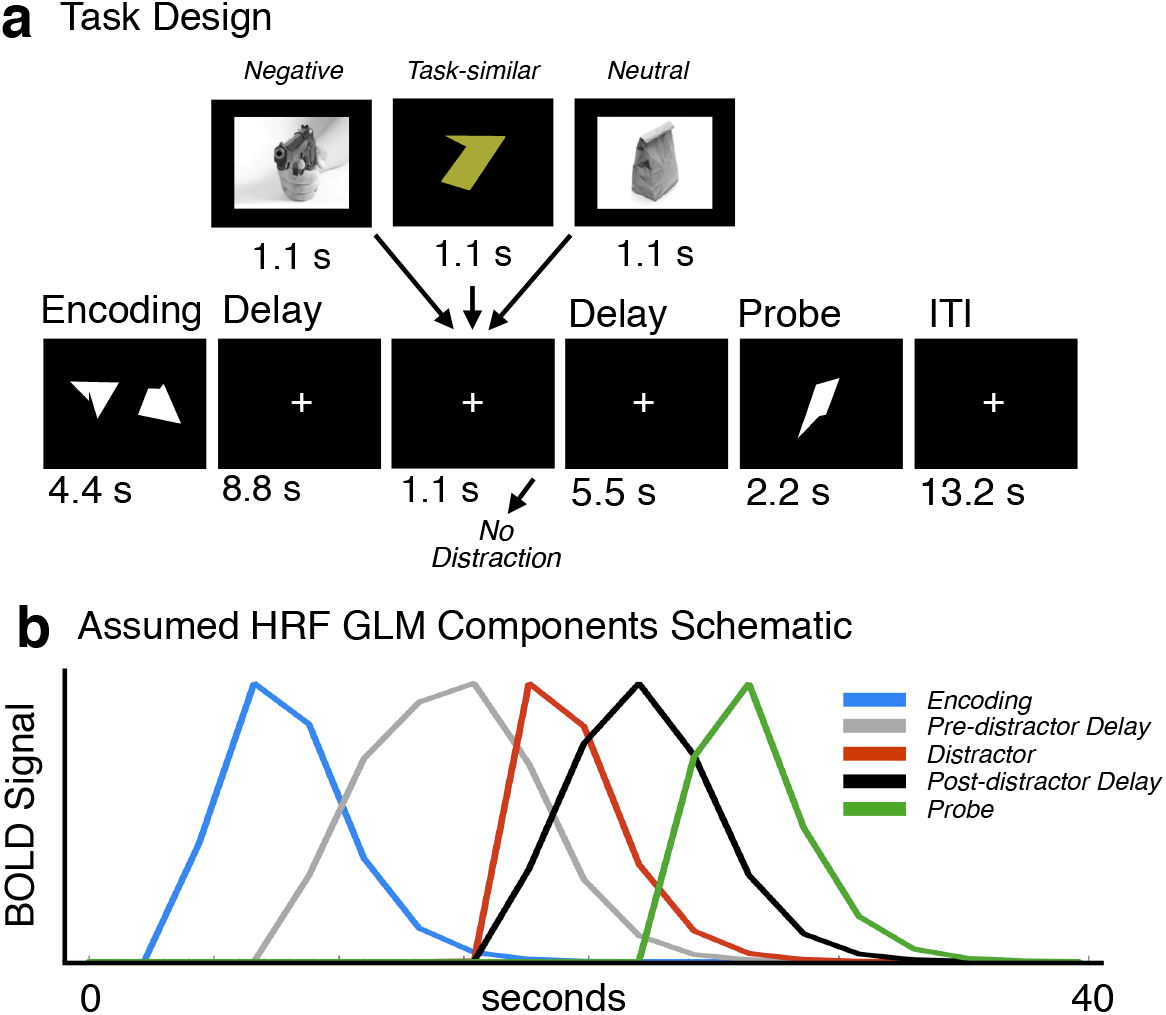
Task design and neuroimaging model schematic. (**a**) Working memory interference paradigm. The task design is displayed along with different trial components and their onsets marked along the timeline. Each box represents a trial component with the duration marked below. Memory sets were presented centrally, subtending a visual angle of 15.75 for 4.4 sec, followed by an 8.8-sec delay. The delay was followed by a 1.1-sec presentation of the distractor (if present) and then by a 5.5-sec post-distractor delay and a probe presented for 2.2 sec. Each trial was followed by a 13.2-sec fixation period (inter-trial interval [ITI]) to allow the hemodynamic response to return to baseline, as employed in our prior work. Distractors were: i) affectively negative complex images, ii) a task-similar geometric shapes of a different color distinguishing it from the probe, iii) neutral complex images, or iv) no distraction. As noted, neutral and negative images were matched on relevant visual characteristics (see **Methods**). (**b**) Assumed hemodynamic response function (HRF) components used in the general linear model (GLM). We modeled 5 different components of each trial: i) Encoding phase (blue), ii) Pre-distractor delay phase (gray), iii) Distractor response (red), iv) Post-distractor delay phase (black), and v) Probe response (green). Distractor response and post-distractor delay were modeled separately for each condition type (i.e. neutral, emotional, task-related and distractor-free trials). These assumed HRF estimates were used to derive the group-level maps. Time courses of activity were extracted for visualization only (see **Methods**).

**Figure 2.**
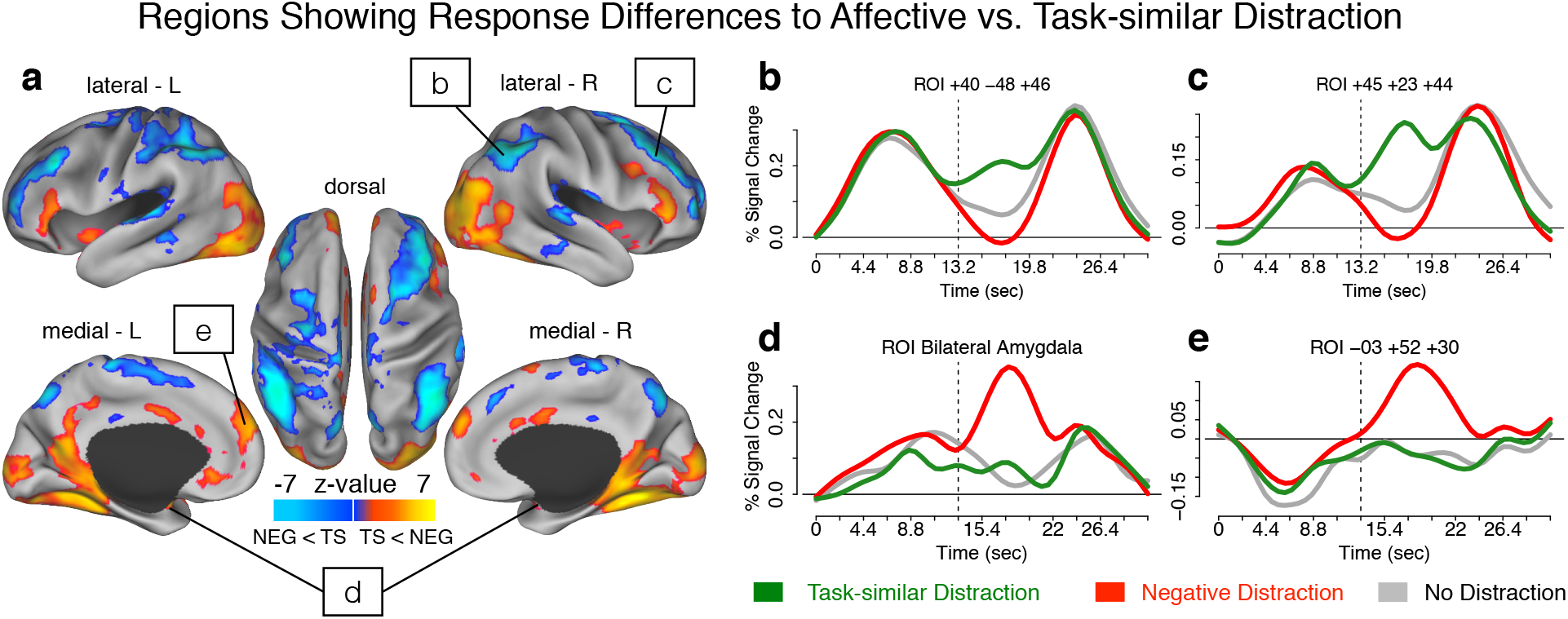
Regions showing dissociable responses to affective vs. task-similar distraction. (**a**) Results from a whole-brain paired t-test directly comparing negative vs. task-similar interference during delayed WM. The map on the left shows regions for which there was significantly higher BOLD signal following negative interference (yellow-orange) and regions for which there was significant higher BOLD signal following task-similar interference (blue). The map highlights brain-wide differences between ventral/dorsal networks in response to opposing distractor types. Of note, these results were obtained using an assumed HRF analysis, specifically modeling distractor onset. The time-courses on the right show un-assumed BOLD signal estimates for a set of exemplar regions, to facilitate visualization: (**b,c**) Effects overlapping with fronto-parietal executive regions typically associated with WM performance, for which there was an increase in BOLD signal following task-similar interference (green time-course), but a drop in BOLD signal following negative interference (red time-course), relative to no distraction (gray time-course). (**d,e**) Effects overlapping with ventral/subcortical regions typically associated with affective/salience responses, for which there was an increase in BOLD signal following negative interference (red time-course), but relatively little or no response to task-similar interference (green time-course), relative to no distraction (gray time-course). The coordinates above each figure highlight the center of mass for a given region (for a full list see **Tables 1-2**). The vertical dashed lines indicate the onset of the distractor.

**Figure 3.**
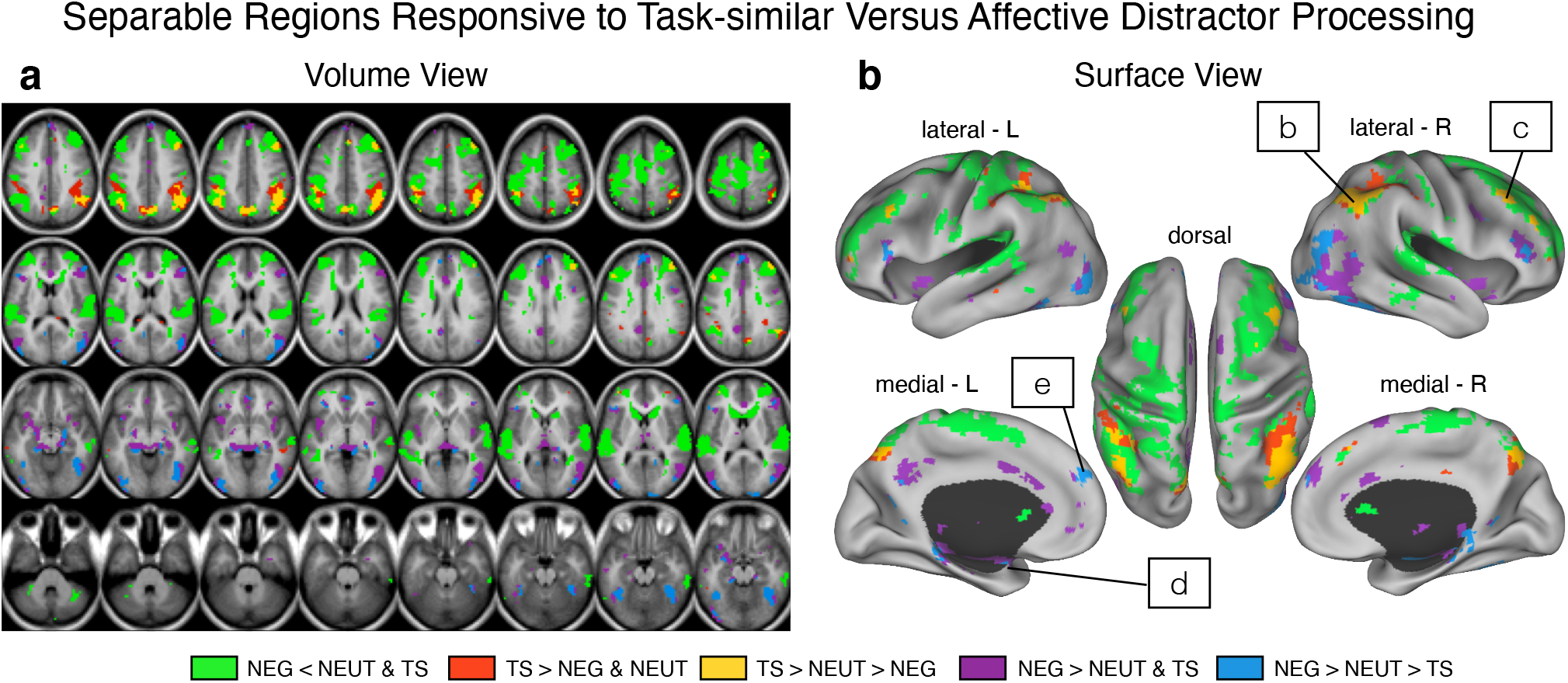
Identification of separable regions responsive to task-similar versus affective distractor processing. The analysis tested for specific and separable response patterns between task-similar distractors (TS) and affectively negative distractors (NEG), using visual neutral distractors as a control (NEUT). Contrasts were computed for each pair of comparisons and formed the following conjunctions to identify dissociable patterns of activity, shown in (**a**) volume and (**b**) surface view: i) where NEG were lower than both NEUT and TS distractor responses (green regions); ii) where TS were higher than both NEUT and NEG distractors (red regions); iii) where TS responses were higher than NEUT, which were also higher than NEG (i.e. a graded response, yellow regions, shown in box b & c); iv) where NEG responses were higher than either TS and NEUT distractor responses (purple regions); and v) where NEG responses were higher than NEUT, which were also higher than TS (i.e. a graded response, blue regions, shown in box e & d). Note that regions identified using this stringent conjunction approach converged closely onto the regions shown in **Figure 2**, confirming separable responses to NEG vs. TS distractors along distinct cortical and subcortical systems. For a comprehensive list of regions that showed TS>NEUT>NEG (yellow) and NEG>NEUT>TS (blue) response patterns see **Table 3**.

For all subjects, the trial sequence was pseudo-randomized with the constraint that no distractor type could appear in more than 3 consecutive trials (to avoid mood induction via negative distractors). The WM memoranda sets were presented centrally subtending a visual angle of 15.75 degrees for a duration of 4.4 seconds followed by an 8.8 second delay. The delay was followed by a 1.1 second presentation of the distractor (if present) and then by a 6.6 second post distractor delay and a probe presented for 2.2 seconds (**Figure 1**). Each trial was followed by a 13.2 second fixation period to allow the neural hemodynamic response to return to baseline. Prior to the start of the experiment each subject was presented with instructions explaining the task and given a brief (8 trial) practice session to demonstrate various trial combinations. During the scanning period visual stimuli were presented through an LCD projector to a screen located behind the scanner, which the subject could see through an angled mirror located above the eyes. The entire experiment was divided into 12 scanning BOLD runs, each lasting 9.2 minutes. There were 180 trials in total with 30 distractor-free trials, 50 negative, 50 neutral and 50 task-similar distractor trials. A subset of participants (24 total) completed a shorter version of the task: 24 distractor-free distractor trials (three 5.09min BOLD runs), 50 negative, 50 neutral and 50 task-similar distractor trials (across six 7.44min BOLD runs) because their data were used to match to a clinical population that could not tolerate a scanning length of 180 trials (see (Anticevic et al., 2011)). We verified that all reported group-level effects held for both cohorts of participants (Anticevic et al., 2011).

### Data Acquisition

Data were acquired on a 3T Tim TRIO Siemens scanner at Washington University Medical School. Functional images were acquired using an asymmetric spin-echo, echo-planar sequence, maximally sensitive to blood oxygenation level-dependent (BOLD) contrast (T2*) (repetition time [TR] = 2200ms, echo time [TE]=27ms, field of view [FOV]=256mm, flip=90°, voxel size=4mm^3^). Each BOLD run contained 251 volumes comprised of 32 oblique axial images, which were acquired parallel to the anterior-posterior commissure. All structural images were acquired using a sagittal MP-RAGE 3D T1-weighted sequence (TR=2400ms, TE=3.16ms, flip=8°; voxel size=1mm^3^).

### fMRI Preprocessing

Functional magnetic resonance imaging data preprocessing steps included: i) Compensation for slice-dependent time shifts; ii) Removal of first 5 images from each run during which BOLD signal was allowed to reach steady state; iii) Elimination of odd/even slice intensity differences due to interpolated acquisition; iv) Realignment of data acquired in each subject within and across runs to compensate for rigid body motion (Ojemann et al., 1997); v) Intensity normalization to a whole brain mode value of 1,000 but without bias or gain field correction; vi) Registration of the 3D structural volume (T1) to the atlas representative template based on 12 normal subjects represented in the Talairach coordinate system (Talairach and Tournoux, 1988) using a 12-parameter affine transform and re-sampled to 1mm^3^ representation (Buckner et al., 2004; Ojemann et al., 1997); vii) Co-registration of the 3D fMRI volume to the structural image and transformation to atlas space using a single affine 12-parameter transform that included a re-sampling to a 3mm^3^ representation; viii) Spatial smoothing using a 6mm full-width at half maximum (FWHM) Gaussian filter.

### fMRI Analysis

We used a general linear model (GLM) approach to estimate task-related activity in each voxel for each subject. Analyses were conducted using the Washington University in Saint Louis Neuro-Imaging Laboratory (NIL) pipelines. First, we employed an assumed response GLM approach to specifically isolate distractor-related activation in addition to other trial components. The model estimated 5 different components of each trial (i.e. encoding, pre-distractor delay, distractor response, post-distractor delay and probe, see **Figure 1b**), obtained by convolving a block function reflecting the neuronal response with a Boynton assumed BOLD response function (Boynton et al., 1996b). Distractor response and post-distractor delay were modeled separately for each condition type, whereas encoding and pre-distractor delay components were not, given that subjects had no advance knowledge of upcoming distractor type. This yielded a total of 11 GLM beta estimates. Of note, given the temporal proximity of the distractor and post-distractor delay components we did not make explicit comparisons between the two components and we specifically used the distractor component for the remainder of analyses. All subsequent statistical analyses used the beta estimates from the *assumed response* GLM model. That is, the assumed response GLM was explicitly used to derive beta estimates of distractor-related signal that were entered into 2^nd^ level analyses, which were used to derive the main-effect maps. In turn, we computed another GLM without assuming a hemodynamic response function (HRF) shape (Ollinger et al., 2001). The un-assumed response was modeled explicitly to provide time course of activity that could be used for qualitative visualization of event-related response patterns. Specifically, for the un-assumed HRF model, the first 15 frames of each trial were modeled and the resulting beta estimates were subsequently extracted for each time point to provide time courses of activity in a given identified region for each condition (see **Figure 2** time courses for an example).

As noted, only the resulting assumed-response HRF beta estimates for task-similar and negative distractors were carried into the 2^nd^ level analysis treating subjects as a random factor. At the 2^nd^-level we computed a two-tailed paired t-test contrasting negative vs. task-similar distractor conditions for all voxels. If the two distractors have a unique impact on dorsal-fronto-parietal vs. ventral-affective activity during WM maintenance, then the statistical map should reveal differences in task-evoked signal patterns between the two distractors. To test this, we applied a stringent whole-brain type I error correction (Z>3 & cluster size >20 contiguous voxels) to the resulting map, ensuring that we capture robust whole-brain effects (see **Tables 1-2**). Next, to isolate specific cortical/subcortical ROIs within the type-I error corrected Z-map, we employed an automated peak-searching algorithm, delineating separate ROIs if they were more than 10 mm apart. These ROIs were limited to no more than 80 mm^3^, in order to preclude creating ROIs that spanned several functionally distinct cortical regions, as done previously (Kerr et al., 2004; Michelon et al., 2003) (see **Tables 1-2** for a comprehensive list of all identified peaks). As noted above, the key objective here was to examine the dissociation, at the whole-brain level, between task-similar and affectively negative interference during WM. While our whole-brain corrected map revealed robust signal differences around the amygdalae, given the *a priori* focus on affective regions, we also verified its location via individually specific anatomically defined amygdala masks. Specifically, as done before (Anticevic et al., 2012c), we used an anatomical amygdala ROI mask based on the current sample, isolated using an automated subcortical segmentation process available through FreeSurfer (Fischl et al., 2002; Fischl et al., 2004). We then applied this bilateral amygdala mask to the statistical map described above to isolate above-threshold voxels specifically within the subject-specific anatomically defined amygdala regions, ensuring anatomical precision.

**Table 1.** Region Coordinates - Negative > Task-similar Distraction.

| X | Y | Z | Hemisphere | Landmark | Max Z Value | Cluster size (mm3) |
| --- | --- | --- | --- | --- | --- | --- |
| -31 | -75 | -12 | left | fusiform gyrus (BA 19) | 7.92 | 10125 |
| 31 | -46 | -14 | right | fusiform gyrus (BA 37) | 7.59 | 12636 |
| -31 | -51 | -12 | left | declive | 7.38 | 8181 |
| 29 | -91 | 20 | right | middle occipital gyrus (BA 19) | 7.18 | 6237 |
| 41 | -82 | 11 | right | middle occipital gyrus (BA 19) | 7.08 | 8370 |
| 37 | -77 | -9 | right | fusiform gyrus (BA 19) | 6.89 | 12690 |
| 15 | -35 | -5 | right | parahippocampal gyrus (BA 30) | 6.58 | 8586 |
| -31 | -93 | 11 | left | middle occipital gyrus (BA 18) | 6.38 | 7209 |
| -11 | -55 | 5 | left | lingual gyrus (BA 18) | 6.40 | 6561 |
| -12 | -101 | 0 | left | cuneus (BA 18) | 6.29 | 5508 |
| 14 | -99 | 8 | right | cuneus (BA 18) | 6.25 | 6075 |
| -18 | -37 | -7 | left | parahippocampal gyrus (BA 30) | 6.15 | 8910 |
| -3 | 52 | 30 | left | superior frontal gyrus (BA 9) | 5.90 | 8424 |
| -36 | 29 | -2 | left | inferior frontal gyrus (BA 47) | 6.00 | 2943 |
| 1 | -53 | -39 | right | cerebellar tonsil | 5.64 | 1836 |
| 21 | -15 | -12 | right | parahippocampal gyrus (BA 28) | 5.60 | 3321 |
| -20 | -16 | -11 | left | parahippocampal gyrus (BA 28) | 5.38 | 1620 |
| -39 | 11 | -12 | left | sub-lobar, extra-nuclear (BA 13) | 5.35 | 2403 |
| -26 | -2 | -15 | left | parahippocampal gyrus, amygdala | 5.38 | 1728 |
| -44 | -85 | -2 | left | inferior occipital gyrus (BA 18) | 5.81 | 4104 |
| 38 | 32 | -1 | right | inferior frontal gyrus (BA 47) | 5.10 | 2565 |
| 52 | -65 | 4 | right | middle temporal gyrus (BA 37) | 5.86 | 6345 |
| 49 | 30 | 16 | right | inferior frontal gyrus (BA 46) | 5.04 | 4239 |
| 13 | -55 | 9 | right | posterior cingulate (BA 30) | 5.01 | 5130 |
| -46 | -76 | 18 | left | middle temporal gyrus (BA 19) | 6.02 | 4860 |
| -48 | 27 | 9 | left | inferior frontal gyrus (BA 45) | 4.77 | 3240 |
| 3 | 19 | 23 | right | anterior cingulate (BA 24) | 4.71 | 2160 |
| 0 | 0 | 33 | left | cingulate gyrus (BA 24) | 4.70 | 2457 |
| -4 | -86 | -3 | left | lingual gyrus (BA 18) | 5.04 | 5292 |
| 13 | -86 | -7 | right | lingual gyrus (BA 18) | 5.56 | 6777 |
| 3 | 43 | -4 | right | anterior cingulate (BA 32) | 4.54 | 1755 |
| -5 | 28 | -7 | left | anterior cingulate (BA 32) | 4.62 | 1161 |
| -3 | 39 | 12 | left | anterior cingulate (BA 32) | 4.50 | 2538 |
| -36 | -10 | -7 | left | sub-lobar, claustrum | 4.37 | 2106 |
| 6 | 10 | 65 | right | superior frontal gyrus (BA 6) | 4.39 | 1026 |
| 15 | -95 | 27 | right | cuneus (BA 19) | 4.83 | 1188 |
| 39 | -4 | -5 | right | insula (BA 13) | 4.30 | 2214 |
| 40 | 10 | 28 | right | inferior frontal gyrus (BA 9) | 4.30 | 1269 |
| -15 | 45 | 46 | left | superior frontal gyrus (BA 8) | 4.40 | 1944 |
| -4 | -18 | 4 | left | thalamus, medial dorsal nucleus | 4.72 | 1215 |
| -6 | -45 | 34 | left | cingulate gyrus (BA 31) | 4.62 | 1701 |
| -44 | -70 | -19 | left | declive | 6.52 | 3537 |
| 26 | 3 | -16 | right | parahippocampal gyrus (BA 34) | 3.85 | 1107 |
| 37 | 17 | -10 | right | inferior frontal gyrus (BA 47) | 4.03 | 783 |
| -42 | -49 | -26 | left | culmen | 5.69 | 2295 |

**Table 2.**
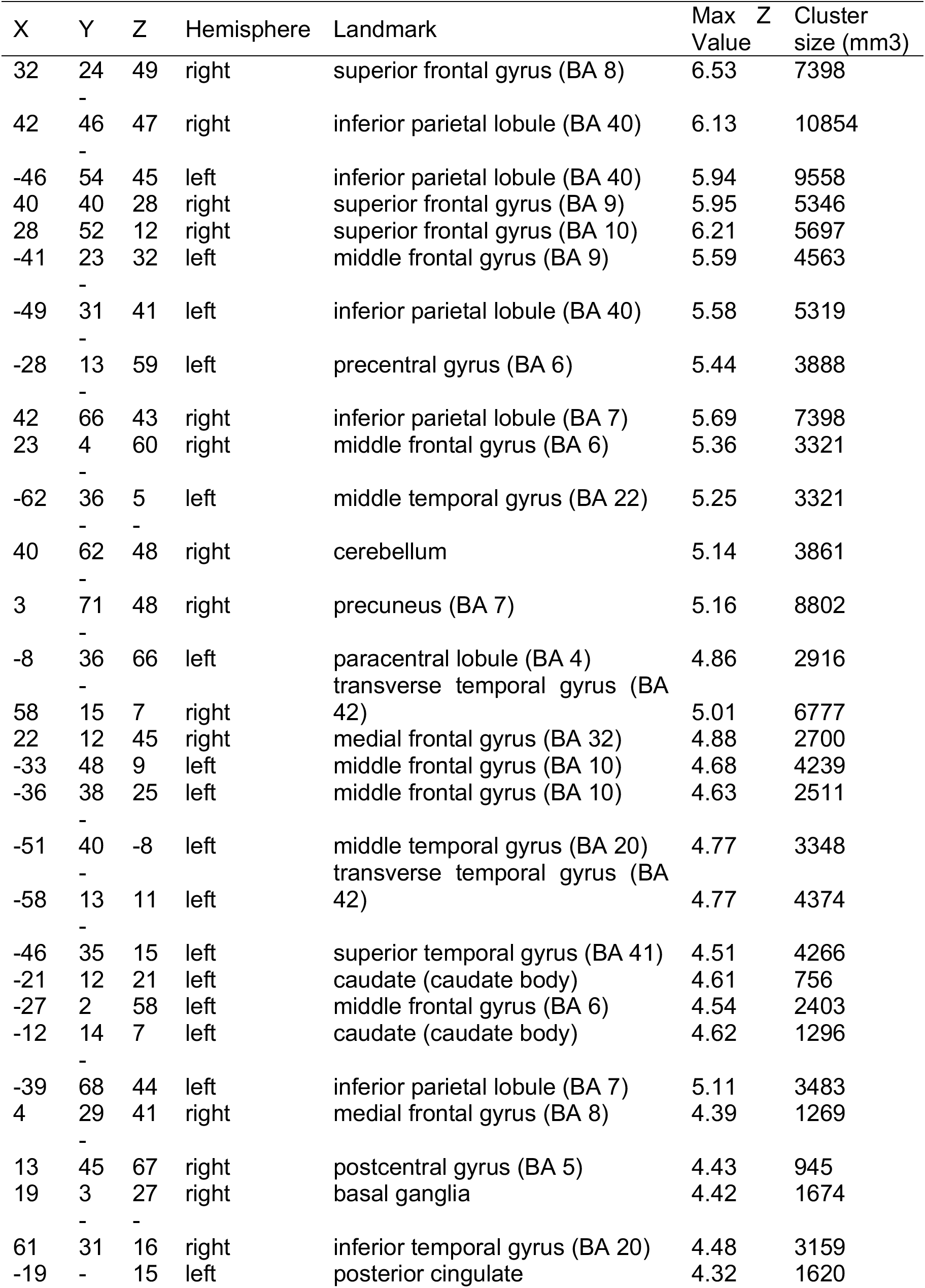
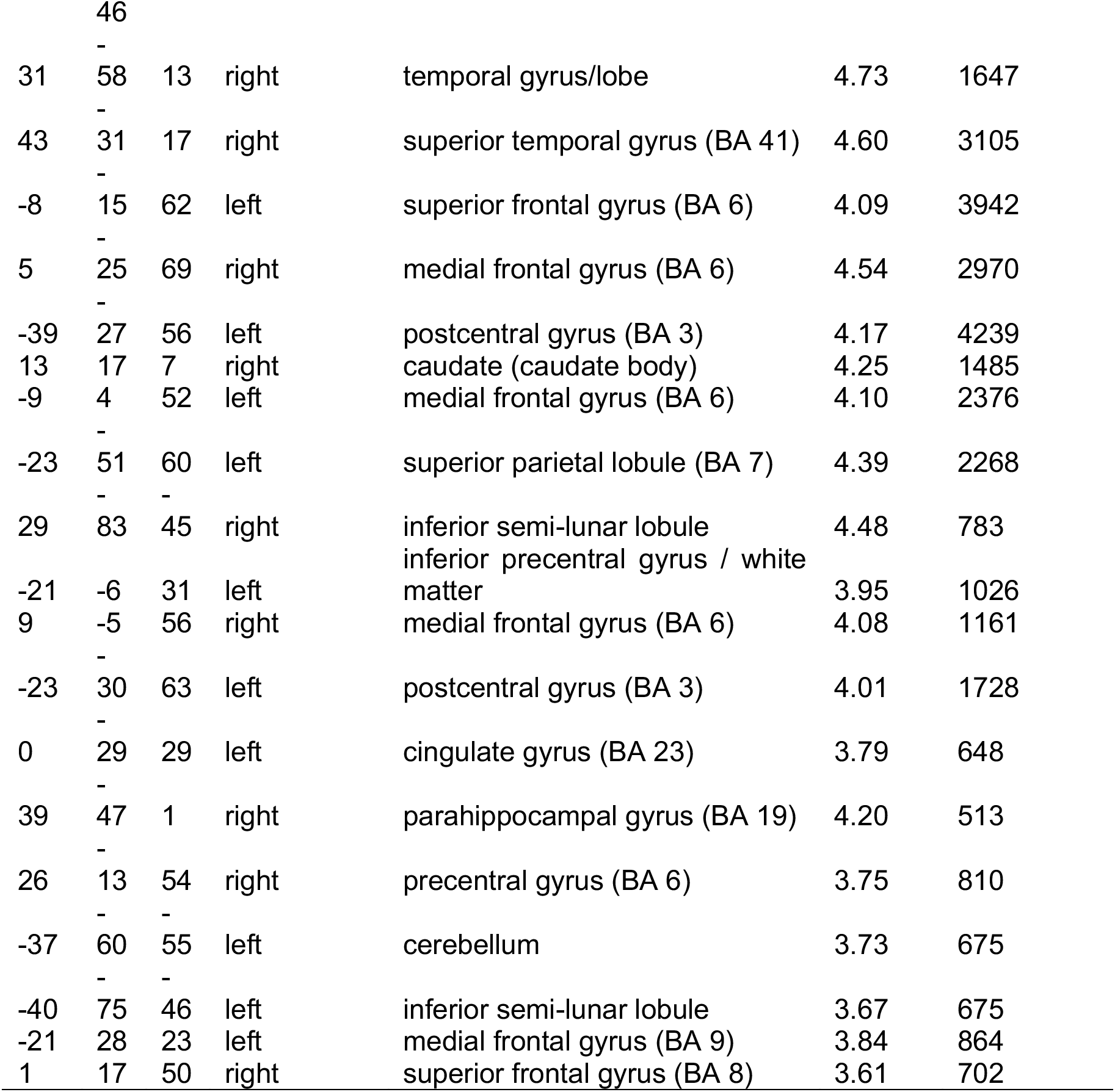
Region Coordinates - Task-similar > Negative Distraction.

### Task-based Functional Connectivity Analyses

Our modeling simulations, which are detailed below, provided insight into how task-based functional connectivity (tb-fcMRI) could be altered in affective vs. cognitive systems as a function of different distraction. To test these connectivity-related predictions, we computed tb-fcMRI analyses for which additional analytic steps were performed:

#### Preprocessing & Analysis

We further preprocessed the BOLD time series via the NIL pipeline to remove sources of spurious variance that can drive between-region functional connectivity: i) High-pass filtering (>0.009Hz) to remove low frequencies and scanner drift. ii) Removal of motion correction parameters, ventricle, deep white matter, and global mean (GMS) signals as well as their first derivatives using the GLM framework. We conducted all subsequent tb-fcMRI analyses on the residual values as done previously (Anticevic et al., 2012d; Anticevic et al., 2010b). Next, we computed the average BOLD signal value for the approximate encoding (time points 3 & 4) and delay period (time points 8 & 9) at each trial for each voxel in the image, as validated in prior studies (Anticevic et al., 2010a; Anticevic et al., 2010b). We averaged two time-points to reduce variability due to possible outlier frames. Next, we concatenated the values into 4-D (brain volume x trial) time series that represented trial-to-trial variability. Extracting only specific time-locked components of the time series, as demonstrated in prior work (Anticevic et al., 2010a; Anticevic et al., 2011; Anticevic et al., 2010b), ensured that the correlations are driven primarily by trial-to-trial variability and not overall task response.

#### Network Definition and Analysis

Our hypotheses focused on the relationship between the fronto-parietal (cognitive) and the bottom-up ventral (affective) networks (although some areas responsive to affective distractors were cortical). To define our networks, while controlling for individual anatomical variability, we used three steps: i) We employed our task-based results as a guide whereby we first isolated all the activation peaks for the main effect map defined above. These included both cortical and subcortical regions in both the cognitive and affective networks. The peak coordinates for these regions are shown in **Table 1** and **Table 2** respectively. ii) Next, we created spherical ROIs (9mm in diameter) in standard space centered on the peak coordinates for each activation cluster, as done previously (Repovs et al., 2011). iii) We masked the resulting ROIs with the individual subject-derived FreeSurfer segmentation of the high-resolution structural image that was registered to the same standard space (Talairach and Tournoux, 1988). This way we excluded any voxels within the group-defined ROIs that did not represent the relevant gray matter for a given individual subject, ensuring we captured individual-specific gray matter signal. While not fully independent of the main task effects, this approach effectively yielded a network definition for tb-fcMRI that captured the areas driven by the task that intersected with subject-specific gray matter.

Next, we extracted the time series for each of these ROIs and computed the ROI-to-ROI correlation matrix across all ROIs for each participant for cognitive-affective network pairs at the encoding and distraction phase of the trial. All obtained correlations for each subject were converted to Fisher-Z (Fz) values. Given no *a priori* motivation to focus on any one specific ROI-to-ROI connection, as done previously (Anticevic et al., 2010b), we averaged Fz values across all connections between the nodes of two networks of interest to produce a single ‘mean Fz’ index of within and between network connectivity. Using this mean Fz index as the dependent measure we computed 2^nd^-level analyses: a 2-way repeated measures ANOVA with *Task Condition* (task-similar vs. affective) × *Task Phase* (encoding vs. distraction) as factors. Here we used *Task Phase* as a factor, given the expectation that at encoding there should be no effect of distraction (as no distraction occurred yet, therefore the encoding phase serves as a control condition in this case).

### Computational Modeling

Presented neuroimaging findings provide neural system-level evidence for the effects of different distractor types on cognition. To further relate these BOLD effects to possible synaptic mechanisms underlying large-scale neural system interactions, we constructed a parsimonious yet biophysically-grounded computational WM model to understand these effects. We specifically implemented interactions between two distinct modules: a task-activated network (denoted as the “cognitive module”) that responds to task-relevant stimuli and performs WM-related computations; and an affective salience network (denoted as the “affective module”) that is activated by task-irrelevant affectively salient stimuli. The two modules interact via long-range projections that implement a functional antagonism. All the presented simulations are built on prior well-validated biophysically-based models of WM, which are spiking local circuit models capable of WM and decision-making computations (Compte et al., 2000; Murray et al., 2014; Wang, 2002). Moreover, the implemented extension to the system-level neural architecture is based on published approaches (Anticevic et al., 2012b), but expanded here to capture interactions between putative ‘affective’ and ‘cognitive’ modules. More specifically, each module is a local circuit of spiking excitatory (E) and inhibitory (I) cells. E-cells interact with one another through horizontal connections mediating recurrent excitation via NMDA receptors and a pool of I cells that mediate feedback synaptic inhibition via GABA receptors. The cognitive module is comprised of two sub-networks: sensory and mnemonic. The sensory sub-module receives task-similar stimulus input.

#### Model Details

For all circuits, we adapted parameters from a previous spatial WM model (Murray et al., 2014). Here we note changes to parameters from that original set. For the sensory and mnemonic modules, pyramidal cells are tuned to angular location on a circle (0-360°) with uniform distribution of preferred angles. The network structure follows a columnar architecture, such that pyramidal cells with similar stimulus selectivity are preferentially connected to each other. To produce differences in WM-related persistent activity across circuits, we modified the height of the recurrent E-E connectivity profile (J_+_) (Compte et al., 2000). Specifically, the synaptic conductance from neuron *j* onto neuron *i* (g_ji_) is scaled by the Gaussian profile:

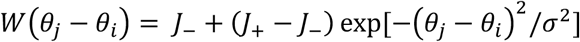

where J_−_is set by normalization of the profile. The sensory sub-module has connectivity profile set by J_+_ = 2 so that it does not support tuned persistent activity in the absence of a stimulus. The mnemonic sub-module, which does support WM-related persistent activity, is set by J_+_ = 3, as in prior work (Murray et al., 2014). For parsimony, we set the affective module to have homogenous connectivity, J_+_ = 1. This is because we do not model selectivity for a particular affective stimulus within this network, and thus it is activated broadly by affectively salient stimuli. The sensory, mnemonic and affective modules each contain N_E_ = 2,048 pyramidal cells and N_I_ = 512 interneurons.

#### Interactions between modules

The model contains projections between modules, originating from pyramidal cells and targeting both pyramidal cells and interneurons. Projections targeting pyramidal cells may be structured based on similarity of stimulus selectivity, and projections targeting interneurons are unstructured and uniform. The sensory module provides structured feed-forward excitation and inhibition to the mnemonic module to relay the task-related stimulus information. This projection is set with g_E-E_ = 600/N_E_ nS, J_+_ = 1.555, σ_E-E_ = 9°, and g_E-I_ = 450/N_E_ nS. The mnemonic sub-module and the affective module send reciprocal long-range projections that are net inhibitory and unstructured (J_+_ = 1). For projections from the mnemonic sub-module to the affective module, g_E-E_ = g_E-I_ = 200/N_E_ nS. For projections from affective module to the mnemonic sub-module, g_E-E_ = g_E-I_ = 10/N_E_ nS. All other inter-module connections are set to zero. Projections between the mnemonic and affective modules create a net inhibitory impact on principal cells, implementing an antagonistic interaction between the two modules.

#### Stimulus

Stimulus input is a 1-sec current pulse to the E-cells in the sensory sub-module with maximum of 0.285 nA and Gaussian profile width of 14.4°. The cue duration was 4.4 sec and the distractor duration was 1.1 sec, to match the experimental protocol (Anticevic et al., 2010a). The model assumes differential routing of task-relevant vs. affective stimuli. The cue stimulus activates pyramidal cells in the sensory network of the cognitive module, which in turn activate the mnemonic network. Task-similar distractors were modeled identically to the initial cues, with same intensity (maximum 0.285 nA) and duration (1-sec current pulse), but with a different stimulus location, such that the distractor appeared at a given angle relative to the original cue. In contrast, affectively salient distractors uniformly activate E-cells in the affective module. We set the negative distractor strength to 0.278 nA for **Figures 4** and **5**.

**Figure 4.**
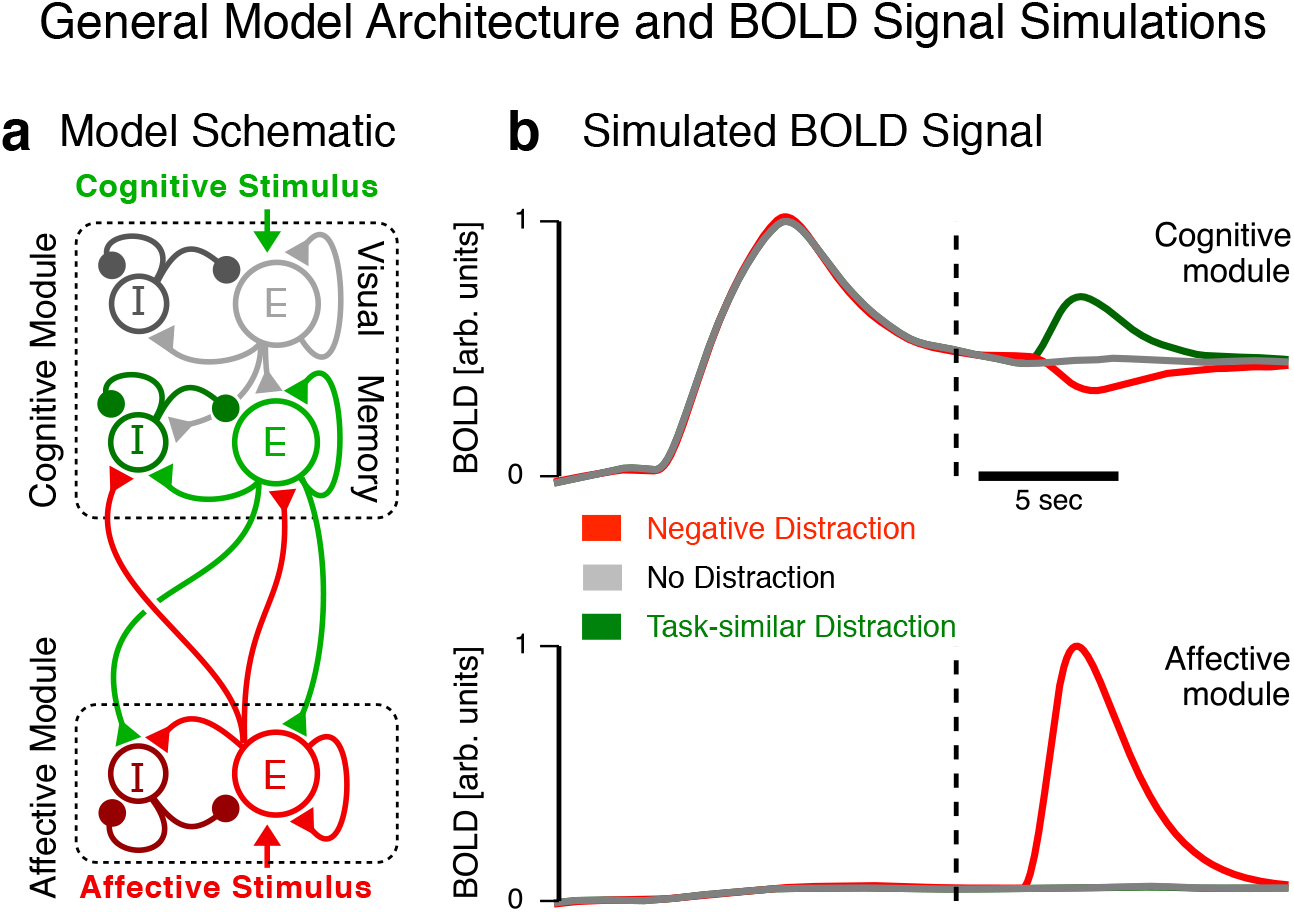
Model scheme and simulated BOLD responses to distractor types. (**a**) The model is comprised of two spiking circuit modules, cognitive and affective, each containing excitatory (E) and inhibitory (I) cells. Task-similar vs. affective stimuli are selectively routed to the cognitive module and affective module, respectively. The cognitive module is characterized by representation and WM-related maintenance of task-similar stimuli. The affective module is characterized by activation by affective stimuli. The cognitive and affective modules interact through long-range projections that are net inhibitory, implementing a functional antagonism. Notably, the cognitive sub-modules do not constitute distinct cortical layers but rather hypothesized neuronal populations with distinct ‘sensory’ vs. ‘mnemonic’ tuning profiles, supported by primate physiology (Ben Hamed et al., 2001; Goldman-Rakic, 1995). (**b**) Simulated BOLD signal qualitatively capture empirical results shown in **Figure 2**. The vertical dashed lines indicate the onset of the distractor.

We quantified the network’s robustness against affective distractors during WM by measuring the distractibility threshold, i.e., the strength of affective distractor input above which the WM signal is lost to suppression of the WM-related activity (**Figure 6**). Specifically, we varied the applied distractor strength (I_app_), and measured the probability of distraction. The distractibility threshold (I_thresh_) was extracted by fitting this relationship with a logistic function: *P(distraction)* = [1 + exp(−(*I_app_* – *I_thresh_*)/*σ*)]^−1^.

**Figure 5.**
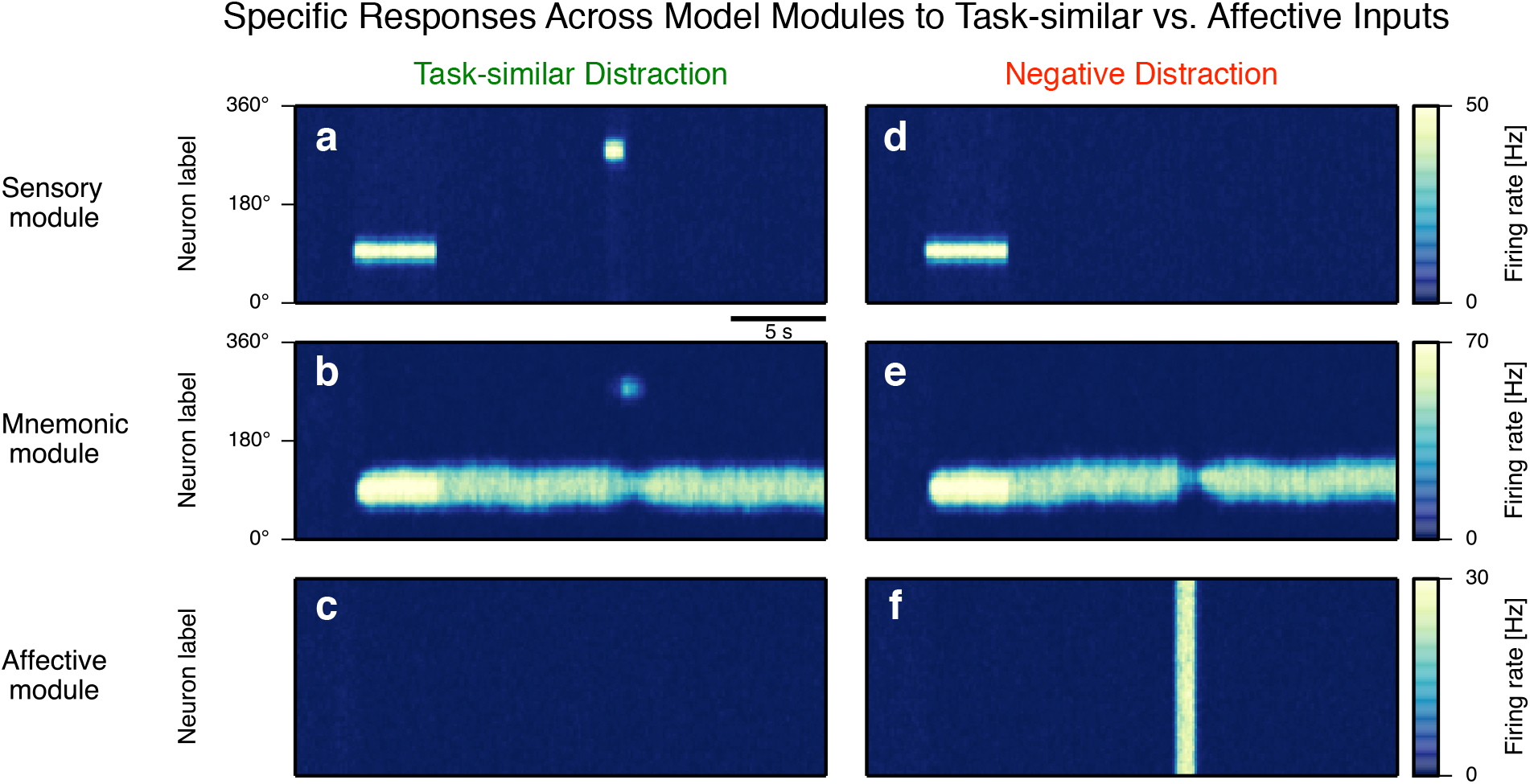
Specific responses for each module to task-similar versus affective stimulus inputs. Spatiotemporal plots of neuronal spiking activity patterns for sensory (**a,d,** top), mnemonic (**b,e,** middle), and affective (**c,f,** bottom) modules in response to a task-similar distractor (left) and an affective distractor (right). (**a,d**) After the initial cue activates the sensory cells, the stimulus identity is encoded by persistent activity within the mnemonic module (panels **b,e**). (Left) A task-similar distractor activates sensory cells (panel **a**), driving competition within the mnemonic module (panel **b**), but no signal is evident in the affective module in this case because the signal was routed to the cognitive module (panel **c**). (Right) A negative distractor activates the affective module (panel **f**), transiently suppressing persistent activity in the mnemonic module (panel **e**), but no signal is evident in the sensory cells in the cognitive module because the signal was routed to the affective module.

**Figure 6.**
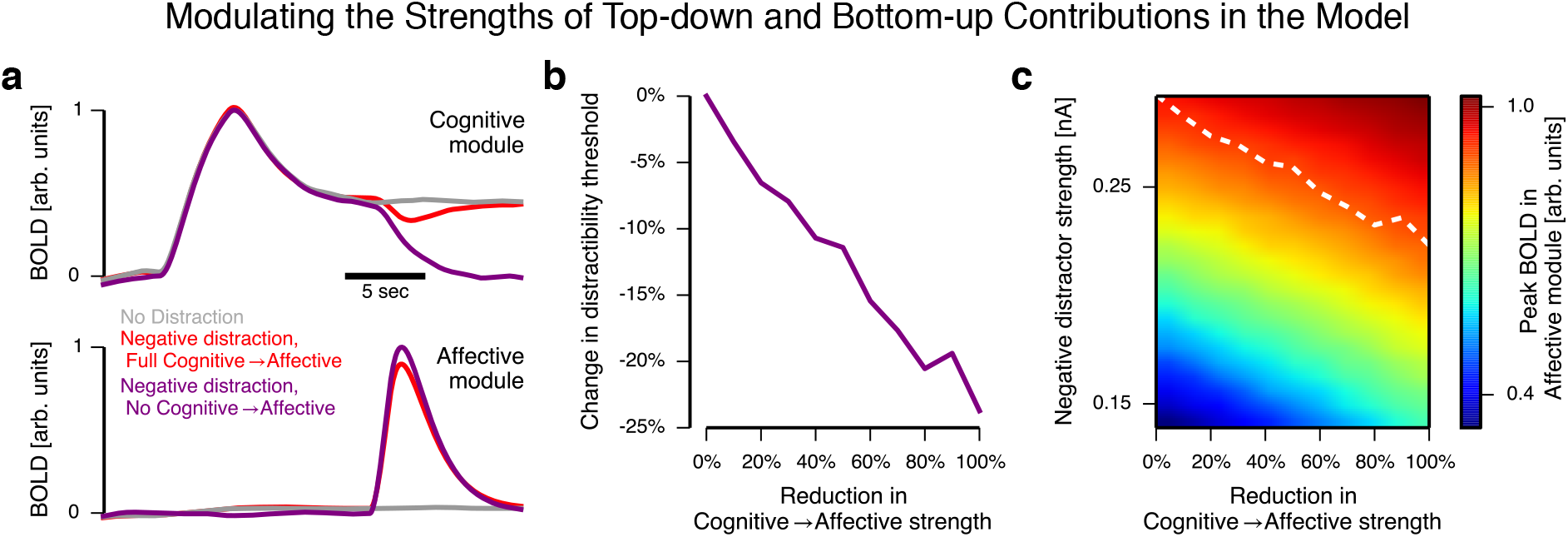
Modulating the strengths of top-down and bottom-up contributions. These simulations examined the role of top-down inhibition from the cognitive module to the affective module. Strength of this projection regulates the strength of activation in the affective module by affective stimuli and the threshold at which an affective distractor can impair behavior. (**a**) BOLD traces for putative affective distraction, with and without top-down projections from the cognitive module to the affective module. Removal of top-down projections disinhibits activity in response to the affective distractor. In the traces shown, the disinhibited distractor response fully collapses the WM delay activity in the cognitive module, leading to a behavioral error in WM. (**b**) Reducing the strength top-down projections decreases the distractibility threshold, above which affective distractors can disrupt WM performance. (**c**) Peak BOLD activation of the affective module in response to an affective distractor. Reducing the strength of top-down projections increases activation of the affective module in response to affective distractors. The white dashed line marks the distractibility threshold in the model simulations.

Simulations were implemented with the Brian neural simulator (Goodman and Brette, 2009). Simulation code is available from the authors upon request.

#### Simulated BOLD Signal

To further relate our modeling findings to observed BOLD results we linked neuronal activity to neuronal ensemble activity to simulated BOLD response. To simulate an approximate BOLD signal in the model, we followed a two-step approach validated in previous studies (Anticevic et al., 2012b; Deco et al., 2004; Stemme et al., 2005): (i) simulate the local field potential (LFP) from synaptic activity in the network; and (ii) convolve the LFP signal with a hemodynamic response function, building on the correlation between LFP and BOLD signals (Logothetis et al., 2001). LFP is calculated as the absolute sum of all non-leak currents (AMPA, NMDA, GABA, and applied external) averaged across all pyramidal cells in a module (Mazzoni et al., 2008). This model of LFP has been used successfully to link spiking circuit models to experimental LFP recordings (Mazzoni et al., 2008). The BOLD signal was then calculated by convolving the LFP signal with a single gamma distribution function of the form:

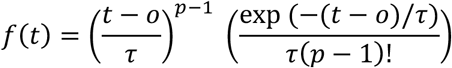

with timescale τ = 1.25 sec, delay *o* = 2.25 sec, and shape parameter *p* = 2. We employed this hemodynamic response function as it was also used to compute the assumed HRF for distinct task phases in the WM trial from the experimental data (Boynton et al., 1996a).

### Data Availability

The data and code that support the findings of this study are available upon request from the corresponding author. A data sharing agreement, project proposal and co-authorship agreement should be submitted.

## RESULTS

### WM Accuracy

We first quantified group-level WM performance across different distractor types. Overall WM accuracy was 78.95%. For the no-distractor condition, WM accuracy was 81.07%. In turn, WM accuracy was 80.41%, 74.99% and 79.33% for task-similar, negative and neutral distractor conditions respectively, consistent with there being a significant effect of distraction across conditions. We verified this by computing a 1-way ANOVA with 4 levels of condition type (task-related, emotional, neutral and distractor-free conditions) [F(3,132)=7.13,p<0.001]. Pair-wise post-hoc t-tests revealed a significant effect of negative distraction relative to no distraction [t(43)=4.27,p<0.00015], verifying that negative distractors exhibited a significant effect on behavior. While numerically lower, other distractor conditions did not significantly statistically differ from the distractor-free condition without distraction on average.

### Task Relevant and Affective Distraction Modulate Distinct Functional Networks

We first tested for dissociable whole-brain effects of distinct distracting information during WM (Anticevic et al., 2010a). As noted, in this experiment, participants performed a modified WM delayed response task (Sternberg, 1969) with three distractor types presented during the delay period: i) affectively negative image (denoted as a “negative distractor”); ii) visually complex neutral image (denoted as a “neutral distractor”); and iii) task-related geometric shape (denoted as a “task-similar distractor”) (**Figure 1**). Prior work established that specific dorsal fronto-parietal areas exhibit signal reductions in response to affective distractors, whereas more ventral and subcortical regions exhibited activations in response to negative distractors. Other more focused studies showed that a similar set of dorsal regions activated in response to task-similar distractors. Yet, it remains unknown if these negative versus task-similar effects differ at the whole-brain level during delayed WM. We hypothesized, based on prior focused regional findings (Dolcos et al., 2008) that affectively negative versus task-similar interference will produce dissociable whole-brain BOLD signal responses during delayed WM.

To test this hypothesis, we first computed a paired t-test using a beta weight representing the assumed HRF BOLD response fit for each distractor type (see **Methods**). Whole-brain results revealed robust differences across dorsal (cognitive) vs. ventral/subcortical (affective) regions for negative vs. task-similar distraction (**Figure 2a**). To visually illustrate this difference, we highlight BOLD responses for exemplar regions using an un-assumed time-course analysis (Ollinger et al., 2001). In two highlighted prefrontal/parietal regions, there was a significantly higher BOLD response to task-similar distractors, but a lower response to negative distractors (**Figure 2b,c**). In contrast, for ventral regions, including the amygdala, we observed a strong response for negative distractors, but a significantly attenuated response for task-similar distractors (**Figure 2d,e**). These results replicate and extend prior regionally focused reports suggesting opposing influence of different distraction on prefrontal signals (Dolcos et al., 2008). These effects also illustrate that task-similar distractors, which match mnemonic features, produce opposing influences on brain-wide neural signals during WM to those observed following affectively negative interference.

### Identification of Separable Responses to Task-similar Versus Affective Distractors

Above we observed significant differences in activity elicited by task-similar versus negative distractors (**Figure 2a**). Next, we sought to establish the specificity of these findings, as these differences can be driven by two possibly distinct properties of the distractor stimuli: affective valence or memoranda task similarity. To further characterize the functional role of identified regions we used neutral stimuli as a comparison (which was omitted in the contrast presented in **Figure 2a** for parsimony). Here neutral stimuli provide a control condition to characterize areas that are uniquely responsitve to affect versus task-similarity. This is possible because neutral distractors differ in one dimension from the other distractor types: i) they differ from task-similar distractors in their similarity to the task cues, but contain no affect; ii) they differ from affective stimili explicity on the affective dimension, but not other visual features on average (see **Methods**). Using this control condition, we tested four possible activity patterns:

First, we hypothesized that regions that are primarly responsive to task-similar inteference would show significant differences between task-similar and neutral distractors, as well as task-similar and negative distractors. However, there should be no significant differences between negative and neutral distractors in such areas (as these regions are not driven by affect). This analysis identified a set of regions that enables sustained maintenance of a memorandum in face of highly similar distractors whose representation might overlap with the representation of the memory set (**Figure 3**, red regions). Second, regions responsive primarily to affective salience should show significant increases in the response to negative stimuli compared to both task similar and neutral stimuli, but no significant differences between the later two (**Figure 3**, purple regions). This contrast would identify areas responsible for processing and orienting to affective distractors specifically (but not other distractor types). Third, regions involved in WM maintenance, but disrupted by affectively salient distractors should show significant decreases in activity specifically after presentation of the negative stimuli in comparison to both neutral and task similar stimuli (Dolcos and McCarthy, 2006). These regions would correspond to a system enabling performance of the ongoing task, which is disrupted when a reorienting affective stimulus appears (Fuster, 1973) (**Figure 3**, green regions). Lastly, we predicted the existence of regions with ovelapping proceses, in which both task-similar and negative distractors elicit significantly different responses than neutral stimuli (**Figure 3**, yellow and blue regions respectively). To identify regions matching these criteria we computed maps of significant differences in brain responses between individual pairs of conditions and then assigned each voxel to one of the above described categories based on patterns of activity.

The results revealed a large set of dorsal regions in which negative distractors significantly decreased WM-related activation compared to both task-similar and neutral distractors (**Figure 3**), replicating prior work (Dolcos and McCarthy, 2006). As predicted, these regions correspond to a subsystem engaged during WM maintenance that is disrupted by affectively salient distractors. Across these regions, a specific subset of areas centered on the fronto-parietal network exhibited graded response, reflecting not only reduced activity due to affective salience, but also increased activity in the case of task-similar distractors (**Figure 3**, foci marked b-c closely corresponding to timecourses shown in **Figure 2b-c**). Indeed, a set of bilateral regions in posterior parietal cortex exhibited only increases in response to task-similar distractors. This revealed regions that are most responsive to distractors similar to the target WM item and are probably directly involved with maintenance of the WM representation. In contrast, a set of ventral regions including occipital cortex, ventral temporal cortex, insula, medial frontal and medial posterior cortex, and ventral PFC showed increased response to affetively salient stimuli (see **Table 3** for a full list of identified ROIs). Across these regions, occipital and inferior temporal regions also show increases to neutral compared to task-similar stimuli, possibly reflecting the difference in the complex content of the stimuli (images vs. geometrical shapes). To summarize, this ‘specificity’ analysis confirmed separable subsystems: i) those that subserve WM task performance with responsiveness to task-similar interference, and ii) those that process affectively salient negative distractor stimuli. This demonstration is critical to establish dissociable cortical responses to task-similarity versus affective salience, motivating the computational modeling aspect.

**Table 3.**
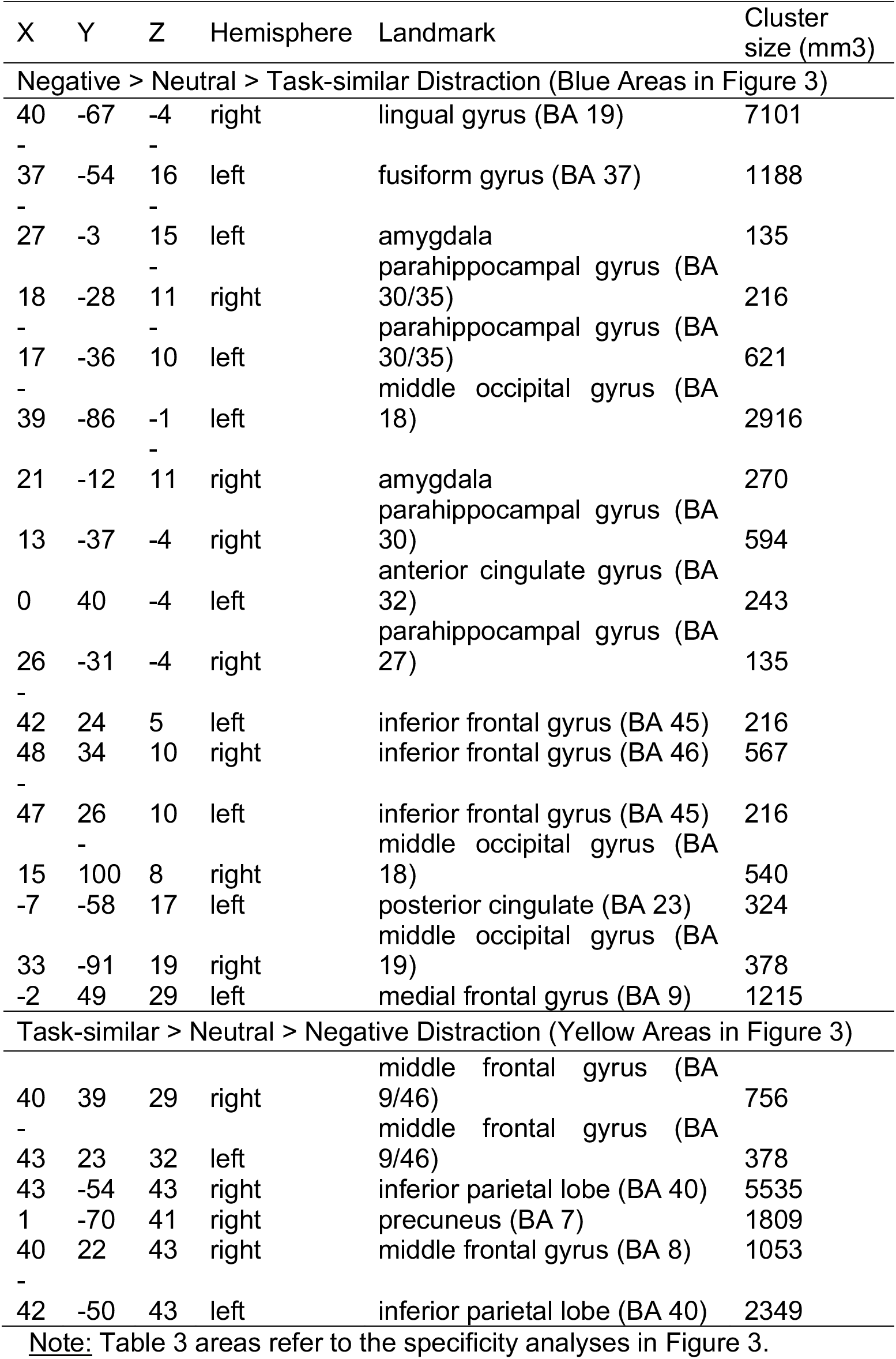
Region Coordinates - Specificity Conjunction Analysis.

### Hypothesized Mechanisms of Affective versus Task-Salient Interface Effects Examined via Computational Modeling

Reported neuroimaging analyses show dissociable effects of task-similar versus negative distractors on WM-related activity across distributed neural networks. These dissociable patterns seem to be confined to distinct systems (**Figure 3**), whereby task-similar distraction increases activity preferentially along fronto-parietal circuits (**Figure 3**, yellow areas). In contrast, affectively salient negative distraction reduces activity in these same areas, while increasing BOLD signal in the medial cortical, ventral and subcortical regions (**Figure 3**, purple areas). Next, we developed a computational neural circuit model to propose a parsimonious mechanism for how such a dissociation might occur. Researchers have previously proposed a putative antagonistic relationship between task-engaged cognitive networks and affective networks (Drevets and Raichle, 1998; Mayberg et al., 1999), yet the synaptic mechanisms of these large-scale neural system interactions remain unknown. To investigate hypothesized mechanisms underlying this functional antagonism on the cortical microcircuit level, which may inform observed BOLD neural system effects, we adapted a well-validated biophysically-based computational model of WM (Compte et al., 2000). We specifically implemented interactions between two distinct modules (**Figure 4a**): a task-activated network (denoted as the “cognitive module”) that responds to task-relevant stimuli and performs WM-related computations; and an affective salience network (denoted as the “affective module”) that is activated by task-irrelevant affectively salient stimuli.

Each module is a spiking circuit of recurrently connected excitatory principal cells and inhibitory interneurons. The cognitive module is comprised of two sub-modules, one sensory and one mnemonic, corresponding to neuronal sub-populations with distinct tuning properties. The sensory sub-module selectively receives structured task-related inputs (cue and task-similar distractor), and represents the task-related stimulus feature in stimulus-selective tuned activity during stimulus presentation, but returns to baseline activity after stimulus withdrawal. Excitatory cells in the sensory sub-module make structured projections onto the mnemonic module. The mnemonic module has stronger structured recurrent excitation, which endows the circuit with the ability to support stimulus-selective persistent activity for WM (Compte et al., 2000; Murray et al., 2014). Thus the cognitive module contains multiple functional cell types that differ in whether they support stimulus-selective persistent activity during WM. This feature is in line with single-neuron recordings from monkey dorsolateral and posterior parietal cortex during memory-guided saccade tasks, in which there are visual cells and delay cells (Ben Hamed et al., 2001; Goldman-Rakic, 1995). The affective module is characterized by a low baseline firing rate but activation at the onset of an affective stimulus, an assumption based on prior neuroimaging studies (Dolcos and McCarthy, 2006). Finally, the microcircuit modules interact through long-range, net inhibitory projections, an architecture suggested by neuroimaging findings (Drevets and Raichle, 1998; Mayberg et al., 1999) and consistent with inter-areal projections onto inhibitory interneurons (Duvarci and Pare, 2014; Medalla et al., 2007a; Timbie and Barbas, 2014). We simulated the BOLD signal for a module by convolving the net synaptic current to pyramidal cells with a hemodynamic response kernel (Anticevic et al., 2012b; Deco et al., 2004) (see **Methods**).

We found that this parsimonious model architecture could closely capture the qualitative pattern of observed empirical BOLD effects identified here and by a number of prior neuroimaging studies (Anticevic et al., 2010a; Anticevic et al., 2011; Dolcos et al., 2008; Dolcos and McCarthy, 2006). To mechanistically capture empirical fMRI observations, the model architecture instantiates two key assumptions that are based on neurophysiological and anatomical findings: differential routing of stimuli according to their task-similar or affective properties, and functional antagonism between cognitive and affective modules. The differential response to distractor types in the cognitive module arises due this routing in combination with the suppressive effect onto the cognitive module induced by the affective module following negative distractors. To enable comparison with empirical observations, we computed a BOLD signal based on model-generated synaptic activity. We found that this model can qualitatively capture the experimentally observed BOLD activity in response to task-similar vs. negative distractors (**Figure 4b**). Specifically, a task-similar distractor increases BOLD in the cognitive module, without strong modulation of the affective module. Conversely, an affective distractor induces an increased BOLD signal in the affective module and a suppression of the BOLD signal in the cognitive module.

Figure 5 shows the differential neuronal processing of distractor types within the circuit modules underlying the BOLD responses. During the initial cue presentation, a tuned subset of sensory cells within the cognitive module is activated, which relays this signal to a subset of mnemonic cells (**Figure 5a**). After the cue has been removed, the mnemonic cells are able to sustain the stimulus-selective pattern of activity throughout the delay, through recurrent excitation and lateral inhibition (Compte et al., 2000) (**Figure 5b**), whereas the affective module remains at a baseline activity level (**Figure 5c**). During presentation of a task-similar distractor, a different subset of sensory cells is activated, which in turn excite corresponding mnemonic cells (**Figure 5a,b**). Competition between cue and distractor representations within the mnemonic sub-module mediates task-similar distractor effects (**Figure 5b**). In the model behavior is compromised when the distractor representation is too strong, and the mnemonic representation switches to represent the distractor (Compte et al., 2000; Murray et al., 2014). In contrast, during presentation of a negative distractor, cells in the affective module are activated (**Figure 5f**). Due to projections from the affective module to the interneurons in the mnemonic module, WM activity is suppressed during negative distraction (**Figure 5e**).

Next we examined the role of projections from the cognitive module to the affective module, which can mediate “top-down” cognitive control of affective processing. We parametrically reduced the overall strength of projections. Because top-down projections are net inhibitory, ongoing activity in the cognitive modules induces suppression of responses in affective module. Weakening top-down projections therefore disinhibits the affective module, increasing its response to the negative distractor. This in turn can disrupt WM activity in the cognitive module, and lead to behavioral errors (**Figure 6a**). Behavioral distraction is characterized by a threshold on the affective stimulus strength, above which the distractor can disrupt WM. We found that weakening top-down strength in the model (i.e. cognitive-to-affective inputs) decreases this distractibility threshold (**Figure 6b**). Weakened top-down strength also smoothly disinhibits the response of the affective module in response to negative distractors (**Figure 6c**). Therefore, net inhibitory projections from cognitive to affective areas provide a hypothesized mechanism whereby ongoing cognitive processing can suppress responses to affective stimuli in ventral cortical and subcortical areas (Ochsner and Gross, 2005; van Dillen et al., 2009).

Importantly, the model architecture makes several predictions. The antagonistic interactions may be evident as anti-correlation between activity patterns in cognitive and affective areas, particularly during processing of negative distractors when affective areas are activated. The model also predicts that errors during negative distraction may be linked to greater responses in affective areas and greater suppression in WM-related areas. Furthermore, the model suggests that one plausible mechanism underlying behavioral errors could be related to increased activation of more ventral areas by affectively salient stimuli. This in turn causes increased suppression of cognitive areas by affective areas, and consequently decreased top-down suppression of affective processing by cognitive areas (i.e. an altered loop).

### Affective Interference Alters Task-based Connectivity Between Networks

Despite its simplified architecture, the modeling results provide an important novel insight that can be extended to the neural system level: affective versus task-similar distraction may uniquely impact the functional connectivity of fronto-parietal (task-similar) versus ventral-affective networks by modulating the activation patterns within each system. That is, if a given distractor type preferentially engages neurons in a given network, then one possible outcome is that the recurrent excitation between areas in a given network will increase. In contrast, affective distraction may preferentially reduce connectivity between networks by increasing the functional antagonism between them, due to a net-inhibitory impact on the task-relevant network by the affective network (**Figure 4**).

We tested this hypothesis via tb-fcMRI, which closely followed prior validated approaches (Anticevic et al., 2012c; Anticevic et al., 2012d; Anticevic et al., 2010b) (see **Methods**). After estimating tb-fcMRI for each subject we computed a within-subject ANOVA with two factors: *Task Phase* (2 levels: encoding and delay/distractor phases) and *Task Condition* (3 levels: no distraction, task-similar and negative distraction). We computed the ANOVA both for the ventral-affective network specifically (**Figure 7a,b**, within-network findings) and between fronto-parietal (task-similar) and ventral-affective networks (**Figure 7c,d**, between-network findings) (for network region selection see **Methods** for details on ROI selection). As noted, the logic for this ANOVA design was to test for a *Task Condition* × *Task Phase* interaction, as there is no expected effect at encoding (since no distraction appears here), but there is a predicted effect following distraction.

**Figure 7.**
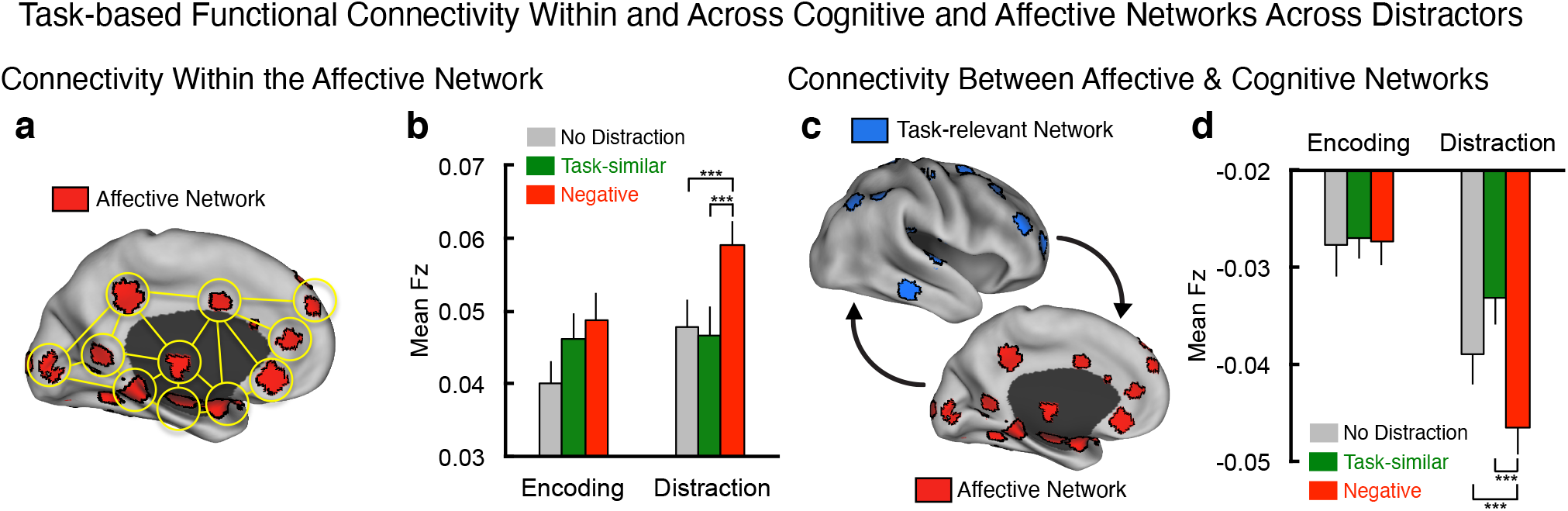
Task-based functional connectivity within and across affective and cognitive networks in response to different distractors. (**a,b**) Task-based functional network connectivity in ‘affective’ areas is significantly increased, specifically by negative distraction during WM delay, in line with the predictions provided by modeling results. (**c,d**) Task-based functional connectivity between areas responsive to task-similar (blue) and affective (red) distractors is significantly decreased, specifically by negative distraction during WM delay, also consistent with the predictions provided by the model suggesting an antagonistic relationship between the modules (see **Figures 4-6**).

Results revealed a significant *Task Condition* × *Task Phase* interaction within the affective/salience network [F(2,88)=5.67, p<0.01] as well as between the affective/salience and task-similar networks [F(2,88)=8.1, p<0.001]. In both cases the effect was driven by a significant connectivity modulation in response to negative distractors during WM delay (**Figure 7b,d**). As predicted, in the case of the affective/salience network, there was a significant increase in tb-fcMRI following negative distraction in comparison to other conditions. Also, in line with predictions, there was a significant reduction in tb-fcMRI across networks following negative interference. Interestingly, there was no significant modulation of tb-fcMRI following task-similar distraction as predicted by the model, either within or across networks (see **Discussion**). Collectively, these effects support the hypothesis that negative interference can significantly modulate functional connectivity both within and across affective/task-similar networks during active cognitive engagement, reflecting the functional antagonism across large-scale neural systems.

### Connectivity Between Affective and Task-similar Networks Predicts Behavioral Performance

The preceding analysis provided evidence consistent with the modeling results, illustrating that the functional connectivity between ventral/affective and dorsal/cognitive networks was reduced as a function of negative interference. This result raised an important secondary hypothesis: an alteration in task-based connectivity may be behaviorally relevant. To test this, we correlated connectivity between affective/task-similar networks with the mean WM accuracy for each subject, specifically for a given distractor condition (**Figure 8**). We found a significant positive relationship between affective/task-similar network connectivity and WM accuracy for the negative condition (r=0.55, p<0.001, 2-tailed; **Figure 8b**). Put differently, those subjects with strongest anti-correlated connectivity between affective/task-similar networks exhibited lowest WM task performance following negative distraction. Such a relationship was not observed for task-similar or distractor-free conditions (r=-0.01 and r=-0.03 respectively, a significant difference in correlation coefficients from those found for the negative condition; **Figure 8d,f**). Here the model results did not necessarily predict that task-similar distraction (or no distraction) would significantly alter the relationship between ventral/affective and dorsal/cognitive areas in relation to behavioral performance, as such distractors would not result in modulation of functional antagonism between these large-scale systems (even though they may affect behavior on some trials). Furthermore, as expected, there was no significant relationship between any of the distracter conditions and WM accuracy during the encoding phase (**Figure 8c,e,g**). These results are consistent with the hypothesis that connectivity alterations between functionally antagonistic affective/task-similar areas are relevant to WM performance specifically following negative distraction: those individuals that have the most functional antagonism following negative affective interference are likely to perform worse.

**Figure 8.**
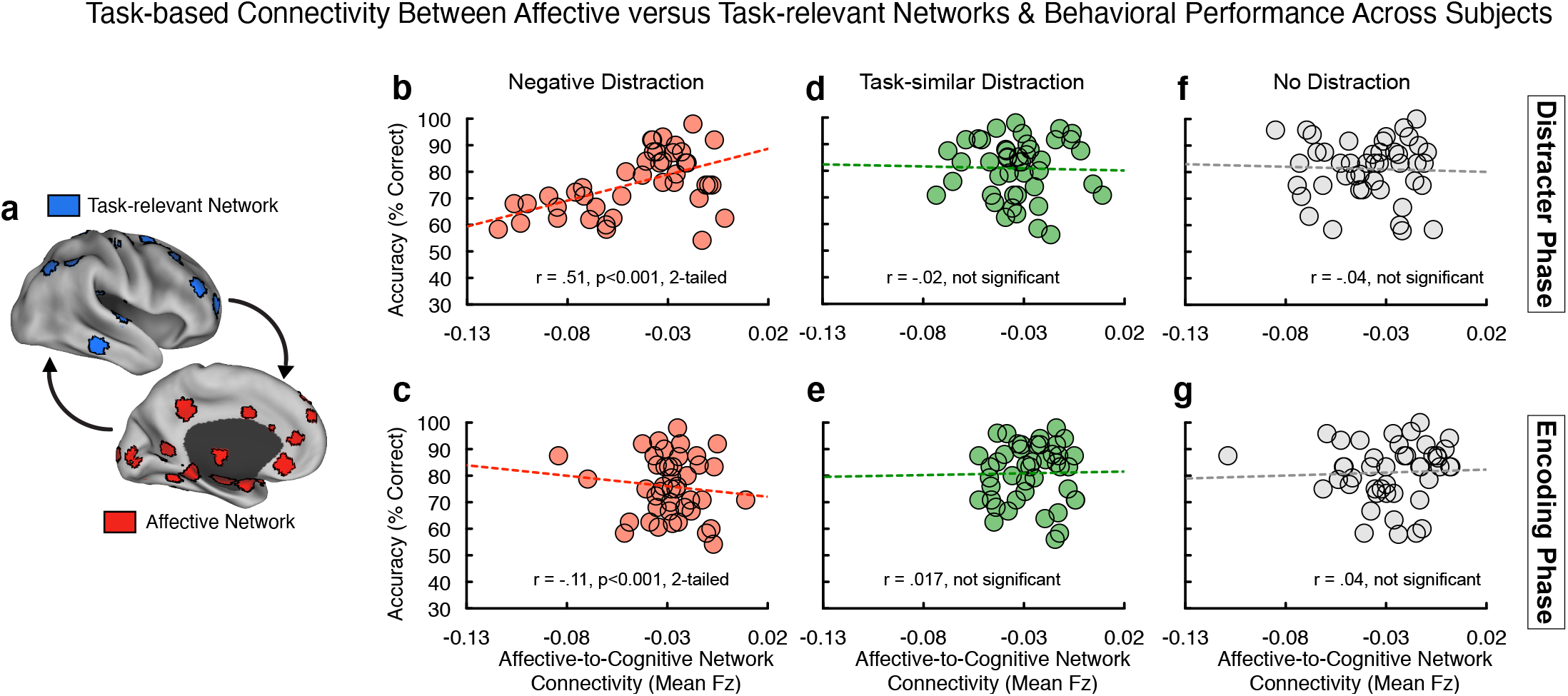
Task-based functional connectivity across ventral/affective and dorsal/cognitive networks predicts behavioral performance during WM. (**a**) We correlated the magnitude of functional connectivity between ventral/affective-salience (red) and dorsal/task-similar (blue) areas for a given condition with behavioral WM performance for that specific condition across subjects. (**b**) Significant positive relationship across subjects (r=0.51, p<0.001, 2-tailed) between task-based networks connectivity and WM performance, specifically for trials containing negative distraction. Of note, we excluded a single outlier case that exhibited below chance performance in the negative distraction condition. The effect remained unchanged when including this case (r=0.55, p<0.001, 2-tailed). (**c**) Notably, there was no such relationship during the encoding condition (r=-0.11, NS). This constitutes a significant difference between correlation coefficients found at the encoding and distractor phases for the negative distractor condition (Z=3.05, p<0.005, 2-tailed). (**d-g**) There was no significant relationship between affective/task-similar connectivity and WM performance for the task-similar distractor condition at either encoding (r=0.017, NS) or maintenance phases (r=-0.02, NS), nor for the WM condition without distraction at either encoding (r=0.04, NS) or maintenance phases (r=-0.04, NS).

### Relationship Between WM Performance and Dissociable Distractor Effects on Dorsal versus Ventral Areas

A related model prediction indicated a differential pattern of activity in dorsal ‘top-down’ areas responsive to task-similar distraction versus ventral/affective areas responsive to negative distraction (see **Figure 6c**). Put differently, negative distraction reduced activity in the ‘cognitive’ module and the magnitude of this reduction was associated with worse WM performance in the model. In turn, higher responsiveness in the ‘affective’ module was associated with higher error rates following affective input, reflecting a putative lack of ‘top-down’ regulation from the cognitive mnemonic module. We tested this model prediction in the following way: i) We identified a subset of subjects that exhibited at least a 10% performance reduction in response to negative distraction relative to their baseline WM performance, to allow for sufficient incorrect trial variability and avoid ceiling WM performance effects (total N=18 subjects). ii) We computed a *Task Condition* (negative vs. task-similar) × *Accuracy* (correct vs. incorrect trials) ANOVA across all the task-similar versus affective-salient regions identified in the primary analysis (**Figure 2**). iii) For the task-similar regions included in this analysis, we computed a conjunction such that selected areas had to overlap with regions that exhibit higher activity for correct WM performance in the absence of distraction. The underlying logic here is that any identified effects of negative distraction had to occur within ‘top-down’ cognitive regions that are explicitly involved in WM performance (as opposed to a wider network that may be responsive to any cognitive task; **Figure 9a**).

**Figure 9.**
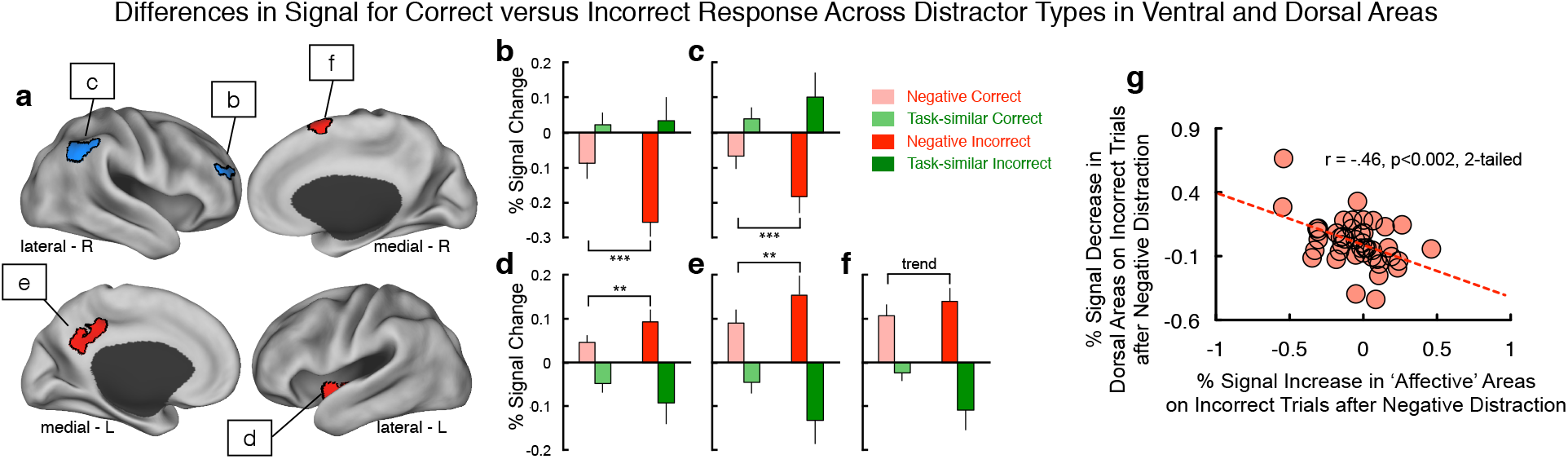
Differences in performance-related signal across cognitive and affective areas in response to different distractor types. (a) We examined the effects of negative versus task-similar distraction on ‘cognitive’ regions (blue) versus ‘affective’ regions (red) as a function of WM performance. We focused on dorsal regions showing main effects of distractor type (see **Figure 2**), which also showed higher WM maintenance activity on correct versus incorrect WM trials (ensuring their involvement in WM task performance). In turn, we examined dissociable effects of task condition and performance across all ‘affective’ areas identified in **Figure 2**, as these regions are not involved in WM maintenance. (**b-f)** We identified 5 total areas, two within the dorsal ‘cognitive’ areas and three within the ‘affective’ areas. All areas showed a double-dissociation between distractor type and WM performance, consistent with model predictions (see **Figure 3**). (**g**) We correlated the percent increase in BOLD signal across the three ‘affective’ areas (x-axis) with the percent reduction in BOLD signal across the two dorsal ‘cognitive’ areas (y-axis) specifically for incorrect WM trials. Effects revealed a significant relationship (r=-.46, p<.002, 2-tailed), suggesting a functional link between the two effects. Notably, we included all cases in this more exploratory ‘individual difference’ analysis to remain maximally powered.

Results revealed two areas in the cognitive regions in which signal following negative distraction was associated with *lower* BOLD signal specifically on incorrect trials. Conversely, we observed the opposite pattern for task-similar distraction, supporting distinct filtering mechanisms (ROI1 (x=33,y=47,z=25), F(1,17)=4.9, p=0.04; ROI2(x=48,y=-53,z=37), F(1,17)=9.43, p=0.007) (**Figure 9b,c**). Within the affective network, we identified the opposite pattern in two cortical regions and one sub-cortical/insular area: responses following negative distraction were associated with *higher* BOLD signal specifically on incorrect trials, whereas the opposite was the case for task-similar distraction (ROI1 (x=-36,y=-10,z=-7), F(1,17)=3.0, p=0.09, trend; ROI2(x=6,y=10,z=65), F(1,17)=6.2, p=0.016; ROI3(x=-6,y=-45,z=34), F(1,17)=3.4, p=0.07, trend) (**Figure 9d-f**). Finally, we tested whether this pattern was associated with individual differences: we correlated the signal increase in ‘affective’ areas associated with incorrect trials with the signal decrease in the ‘cognitive’ dorsal areas associated with incorrect trials, over all cases for maximal power in this more exploratory analysis (**Figure 9g**). Results revealed a significant relationship (r=-.46, p<.002, 2-tailed): subjects with greatest reduction in BOLD signal in ‘cognitive’ dorsal areas exhibited the greatest increase in the ‘affective’ areas following negative distraction, specifically on incorrect WM trials. Collectively, these analyses suggest a dissociable effect of negative interference on ‘top-down’ areas involved in WM versus areas involved in responding to negative distraction.

## DISCUSSION

This study combined functional neuroimaging and computational modeling to provide novel insights into interactions between affect and cognition at the neural system level in humans. We examined a putative ‘antagonistic’ interaction between regions responsive to affective interference versus areas responsive to task-similar distraction during WM – a canonical higher-order cognitive process (Wang, 2001). To generate hypotheses regarding computational principles behind these effects we implemented a circuit-based WM model, extended to the neural system level, generating testable predictions, which can be examined empirically. Collectively, present results offer three insights: i) the presence of large-scale dissociable effects of task-similar interference versus affective interference on WM-related BOLD signal at the whole-brain level, extending prior regional effects (Dolcos et al., 2008). ii) a parsimonious yet biophysically-based computational mechanism for these dissociations based on net inhibitory synaptic-level interactions between neural systems; iii) distinct effects of negative versus task-similar interference on task-based connectivity and their relationship with WM performance, which were predicted by the model architecture and confirmed empirically.

### Dissociable Mechanisms Supporting Task-similar versus Affective Interference

Affective interference has a unique effect on neural activity during cognitive engagement (Ochsner and Gross, 2005; Okon-Singer et al., 2015). This was illustrated during a delayed WM task whereby affective distractors were associated with reduction in WM delay activity (Dolcos and McCarthy, 2006). Subsequent studies identified dissociable effects between task-similar and affective distractors when focused on select regions (Anticevic et al., 2010a; Dolcos et al., 2008). However, a key gap in knowledge relates to whether these effects are brain-wide properties or focal effects. We tested this hypothesis in two ways: First, we identified a robust and brain-wide difference in neural systems responsive to task-similar versus affective interference during WM (**Figure 2**); second, we confirmed the specificity of these effects relative to neutral distractors (**Figure 3**). Both analyses revealed distinct BOLD responses to affective versus task-similar interference, following a ventral/dorsal separation: affective distractors increased signal in affective/ventral areas (with some exceptions in midline cingulate cortical regions). Conversely, task-similar distractors increased signals in dorsal areas involved in WM maintenance.

Similar effects were demonstrated in classic primate physiology WM experiments (Fuster, 1973; Miller et al., 1996): activity of prefrontal neurons during WM in monkeys was selectively attenuated when the animal was presented with a biologically meaningful affective distractor (cry of another monkey). These findings confirm an antagonistic neural architecture between cognitive and affective areas, capable of flexibly engaging distinct incoming distraction and extend prior work in two ways: first, the effects were observed across widespread neural systems supporting a general organizing principle; second, the effects were specific to the distinction between affective and task-relevant interference.

### Computational Mechanisms Underlying Dissociable Distractor Effects

We found that the modeling architecture can closely reproduce BOLD effects across brain areas; task-similar distractors increase BOLD signal in the cognitive module, whereas affective distractors decrease BOLD signal in the cognitive module and increase signal in the affective module, capturing empirical effects. The model architecture is based on findings from physiology and anatomy. One key property of the model architecture is differential routing of stimulus processing according to its task relevance and affective saliency: cue and task-similar distractors are routed to the cognitive module and negative distractors are routed to the affective module. Single-neuron recordings in the monkey dorsolateral prefrontal cortex have shown that distractors are differentially filtered based on their task relevance (Artchakov et al., 2009; Everling et al., 2006). For the ventral affective system, and in particular the amygdala, converging studies have demonstrated preferential activation by affectively salient stimuli, especially those with negative affective valence (Phelps and LeDoux, 2005). This specificity of stimulus routing contributes to the increased activation of the cognitive module by task-similar distraction and activation of the affective module by negative distraction.

In turn, the model implements functional antagonism between the two modules through reciprocal net inhibitory projections, an architecture hypothesized by neuroimaging studies (Drevets and Raichle, 1998; Mayberg et al., 1999). There is anatomical and physiological evidence supporting antagonistic interactions in both directions between cognitive and affective areas. In the model, each projection pathways mediates distinct effects on the processing of negative distraction during cognition. The net inhibitory projection from the affective module to the cognitive module mediates distractibility by negative stimuli (**Figure 6**). This allows for disruption of ongoing cognitive computations, facilitating rapid deployment of attention to novel and survival-relevant stimuli (Ferri et al., 2016; Pessoa and Adolphs, 2010). Suppression interactions may be mediated by long-range projections that target inhibitory interneurons. Projections from amygdala to prefrontal cortex target specific populations of inhibitory neurons (Timbie and Barbas, 2014). Within prefrontal cortex, different long-range cortico-cortical projections, from anterior cingulate cortex versus dorsolateral prefrontal cortex, make distinct patterns of synapses onto excitatory and inhibitory neurons, which could potentially implement the net inhibitory projections (Medalla and Barbas, 2009). Human and rodent studies provide converging evidence for a pathway by which prefrontal cortex may apply top-down inhibitory control of the amygdala via inhibitory interneurons (Duvarci and Pare, 2014; Medalla and Barbas, 2009; Medalla et al., 2007b). The net inhibitory projection from the cognitive module to the affective module mediates a flexible, top-down inhibitory control of affective processing. This model architecture provides enhanced protection of cognitive processing by suppressing responses to incoming affective stimuli (**Figure 6**). Consistent with top-down inhibition of affective systems by the cognitive systems, we observed that cognitive load can reduce the activation of affective areas by negative affective stimuli (van Dillen et al., 2009).

Both long-range projections in the model’s antagonistic architecture contribute to network-level anti-correlation as observed via functional connectivity. The strengths of these net-inhibitory projections thereby have congruent impacts on functional anti-correlation. This is in contrast to their opposing impacts on WM function (**Figure 6**). The projection from the affective module to the cognitive module mediates suppression of the cognitive module by negative distractors. Strengthening this projection lowers the distractibility threshold. In contrast, the projection from the cognitive module to the affective module mediates top-down regulation of affective processing. Strengthening this projection reduces the impact of negative distraction on WM. Therefore, during negative distraction, the bottom-up projection impairs WM performance whereas the top-down projection improves WM performance. Observed functional anti-correlation between networks does not distinguish contributions from these projections (see **Limitations**). However, during the negative distraction, the affective network is strongly activated, potentially augmenting the bottom-up contribution to anti-correlation.

In addition, the modeled reciprocal antagonism across networks may be a common motif for large-scale system interactions in the brain. A similar architecture was used to model another well-established relationship between anti-correlated networks during cognitive tasks – namely the fronto-parietal control system (FPCN) and the default mode network (DMN) (Anticevic et al., 2012b). Understanding how these processes interact and in turn impact behavior is vital to inform the cognitive neuroscience of functional segregation in the human brain, which can have clinical relevance (Anticevic et al., 2012a; Anticevic and Corlett, 2012). Such dynamics of cognition-emotion interactions are relevant for multiple neuropsychiatric illnesses, including major depressive disorder (MDD) and post-traumatic stress disorder (PTSD). In MDD, there is evidence of localized hyper-activity in a cortical area, the subgenual cingulate cortex (Mayberg et al., 1999; Mayberg et al., 2005).

Similar to our findings presented above, hyperactivity in this area could impair its ability to shutoff and suppress cognitive areas, consistent with transcranial-magnetic stimulation findings (Fox et al., 2012). In PTSD, prefrontal-mediated inhibitory control over affective areas may be effectively weakened (Koenigs and Grafman, 2009; Milad and Quirk, 2012; Morey et al., 2009). The model presented here can be used to test how putative circuit abnormalities in these disorders disrupt the balance of antagonistic interactions between cognitive and affective neural systems. Building on these effects, we hypothesize that dysregulation of the functional antagonism between cognitive and affective areas may contribute to cognitive deficits observed in PTSD, anxiety, and depression, possibly via distinct mechanisms that may compromise the appropriate large-scale coordination between neural systems (Anticevic et al., 2012a).

Finally, the model generated predictions for neural activity associated with errors induced by negative versus task-similar distraction. Specifically, the model predicted greater WM error rates in response to more anti-correlated connectivity between the two modules as well as greater error-related signal reductions in the cognitive module, but greater error-related signal increase in the affective module. Both effects were confirmed empirically (**Figures 8-9**). These model-predicted empirical findings illustrate that the identified whole-brain results are behaviorally relevant in terms of functional interactions (i.e. connectivity), performance on a trial-by-trial basis and across subjects. Such performance-relevant effects add additional insight, suggesting that possible breakdowns in the appropriate functional antagonism between cognitive/affective regions may have behavioral consequences for optimal cognitive performance. In turn, this insight can inform understanding of psychiatric conditions where interactions between affective and cognitive processes are compromised (e.g. PTSD, depression, anxiety, and schizophrenia).

### Limitations

The model introduces a parsimonious circuit mechanism, grounded in known neurobiology, that can capture the core empirical findings presented here. However, the minimal, modular architecture limits the extent of the model’s predictions beyond these features. Future modeling studies should extend beyond reduced modules to incorporate complex large-scale networks (Deco et al., 2015). The model represents the ‘affective’ module as a homogeneous pool of neurons, and thereby captures network-level interactions rather than specific computations implemented within affective areas. Furthermore, it remains to be established if this ‘antagonistic’ architecture can flexibly engage with fronto-parietal areas at times when the affective input is actually task-relevant (Spreng et al., 2010). The model is explicitly built around a delayed WM framework. It will be important to extend this approach to test if similar computational mechanisms operate across other cognitive processes (e.g. decision making, (Wang, 2008)). This study is also limited in its capacity to make causal inferences due to the indirect nature of the BOLD signal. As described above, functional connectivity is limited in its ability to resolve distinct antagonistic contributions to the observed anti-correlation, due to the lack of directionality in functional connectivity. Convergent evidence is needed from experimental transcranial magnetic stimulation (Fox et al., 2012) or pharmacological studies (Anticevic et al., 2014) to establish causality by impairing ‘top-down’ or ‘bottom-up’ contributions. Another consideration related to causality in the context of presented functional connectivity analyses is the possibility of "shared task input" issues. Mainly, signals from two regions could be driven by a single task input, and therefore show very similar trial-by-trial fluctuations, but there may be little direct interaction between these two regions. Additional convergent techniques that could be used in future studies may involve partial physiological interaction analysis or the "background connectivity" approach (Al-Aidroos et al., 2011).

## Conclusions

This computational cognitive neuroscience investigation blends neuroimaging and computational modeling in the context of delayed WM faced with distinct distracting information. We extend the WM circuit model to the level of neural systems, to provide three insights to into the ‘antagonistic’ interaction between affect and cognition in the human brain: First, we observed whole-brain dissociable effects of task-similar interference versus affective interference on WM-related BOLD signal. Second, we implemented a parsimonious yet biophysically-based computational mechanism for these dissociations based on net inhibitory synaptic-level interactions between neural systems. Third, the model generated testable predictions following negative interference and its effects on large-scale network connectivity, which were confirmed empirically. These findings highlight an antagonistic architecture between cognitive and affective systems during WM, capable of flexibly engaging distinct distractions during cognition, with implications for psychiatric conditions where the interaction of affect and cognition may be compromised.

## ACKNOWLEDGMENTS

Financial support for this study was provided by NIH grants DP5OD012109-01 (PI: AA), 1U01MH121766 (to AA), R01MH112746 (PI: JDM), 5R01MH112189 (to AA), 5R01MH108590 (PI: AA), NIAAA grant 2P50AA012870-11 (PI: AA), NSF NeuroNex grant 2015276 (PI: JDM), the Brain and Behavior Research Foundation Young Investigator Award (PI: AA), SFARI Pilot Award (PI: JDM & AA), by BlackThorn Therapeutics (PI: JDM & AA). We would like to thank the McDonnell Foundation for providing initial funding for collecting these data. We thank Dr. Deanna Barch for her helpful input and advice during preparation of this manuscript.

